# Androgen deprivation triggers a cytokine signaling switch to induce immune suppression and prostate cancer recurrence

**DOI:** 10.1101/2023.12.01.569685

**Authors:** Kai Sha, Renyuan Zhang, Aerken Maolake, Shalini Singh, Gurkamal Chatta, Kevin H. Eng, Kent L. Nastiuk, John J. Krolewski

## Abstract

Androgen deprivation therapy (ADT) is an effective but not curative treatment for advanced and recurrent prostate cancer (PC). We investigated the mechanisms controlling the response to androgen-deprivation by surgical castration in genetically-engineered mouse models (GEMMs) of PC, using high frequency ultrasound imaging to rigorously measure tumor volume. Castration initially causes almost all tumors to shrink in volume, but many tumors subsequently recur within 5-10 weeks. Blockade of tumor necrosis factor (TNF) signaling a few days in advance of castration surgery, using a TNFR2 ligand trap, prevents regression in a PTEN-deficient GEMM. Following tumor regression, a stem cell-like population within the tumor increases along with TNF protein levels. Tumor cell lines in culture recapitulate these *in vivo* observations, suggesting the possibility that stem cells are the source of TNF. When TNF signaling blockade is administered immediately prior to castration, tumors regress but recurrence is prevented. This implies that a late wave of TNF secretion within the tumor – which coincides with the expression of NFκB regulated genes – drives recurrence. The inhibition of signaling downstream of an NFκB-regulated protein – chemokine C-C motif ligand 2 (CCL2) – prevents post-castration tumor recurrence, phenocopying post-castration (late) TNF signaling blockade. CCL2 was originally identified as a macrophage chemoattractant and indeed at late times after castration gene sets related to chemotaxis and migration are up-regulated. Importantly, enhanced CCL2 signaling during the tumor recurrence phase coincides with an increase in pro-tumorigenic macrophages and a decrease in CD8 T cells, suggesting that recurrence is driven at least in part by tumor immunosuppression. In summary, we demonstrate that a therapy-induced switch in TNF signaling induces an immunosuppressive tumor microenvironment and concomitant tumor recurrence.

## INTRODUCTION

Androgen deprivation therapy (ADT) via surgical castration (1) and currently via medical therapy employing gonadotropin releasing hormone analogues (2), has been an effective treatment for advanced primary and recurrent prostate cancer (PC) for many decades. Within the past several years, the addition of either the microtubule toxin docetaxel or the androgen biosynthesis inhibitor abiraterone to ADT has improved the medical therapy (3). However, these ADT-combination therapies have only marginally extended overall survival and the fundamental issue for patients with advanced primary and recurrent PC remains the same: ADT (or an ADT-combination) rarely cures PC and resistant tumors are eventually lethal. Thus, there is a need to develop additional therapies for PC, preferably by identifying molecular targets that are mechanistically linked to the development of ADT resistance. ADT is effective because the growth of almost every prostate cancer is dependent on a program of gene expression mediated by the androgen receptor and a set of critical co-regulators. The androgen receptor (AR) gene is activated by genetic alteration in only a small fraction of primary PCs (4). Instead, one or two alterations of a small group of other genes (5) is sufficient to re-program the AR cistrome to transcribe a set of genes that promote immortalization and tumorigenesis (6). Additional alterations in a second class of genes driving proliferation and growth – such as the PTEN tumor suppressor or the c-MYC oncogene – completes the process of transformation to prostate cancer (5).

Tissue reconstitution studies (7) have shown that androgen blockade can act indirectly via soluble factors derived from the tumor micro-environment (TME). Consistent with this paradigm, in which proteins are secreted into the TME and act on the cell of origin (autocrine) or on other cells within the TME (paracrine), we have previously investigated the response of prostate cancers to androgen deprivation (8–13), using cell culture and murine models of human prostate cancer. A recurrent theme of this research has been the role of tumor necrosis factor (TNF) as a pleotropic signaling mediator within the PC TME. TNF binds and trimerizes its cognate receptor and signals via a complex network of intercellular protein-protein interactions to promote either an anti-tumor or pro-tumor fate, depending on the cellular context. In this report, we examine the role of TNF and TNF-regulated genes in the response to castration in a genetically engineered mouse model (GEMM) of PC driven by deletion of the PTEN gene (14). Castration induces regression and subsequent recurrence in this GEMM, resembling the onset of castration resistant prostate cancer (CRPC). Both responses are dependent on TNF signaling and can be separately blocked by differentially timing the administration of a TNF ligand trap molecule. At late times post-castration, when the tumor is beginning to recur in a castration resistant form, TNF induces NFκB-mediated gene expression. Moreover, we demonstrate that the NFκB-induced gene CCL2 – which encodes a protein that is chemotactic for immune suppressive cells such as tumor associated macrophages (TAMs) – is necessary for recurrence in the PTEN-loss model as well as in a second GEMM of PC that is driven by MYC over-expression. Thus, a TNF signaling switch induces immunosuppression in the tumor microenvironment to promote castration-resistant tumor recurrence.

## MATERIALS AND METHODS

### Animals and tumor imaging

All animal studies were approved by the Roswell Park Comprehensive Cancer Center (RPCCC) institutional animal care and use committee. Male Pb-Cre4; PTEN^fl/fl^ mice (14) and Hi-MYC (15) were breed in the RPCCC laboratory animal shared resource. Mice were housed in environmentally-controlled conditions on a 12-hour light/dark cycle with water and food. For both genetically engineered mouse models (GEMMs) we studied, male pups were genotyped and tumor volumes were monitored by three-dimensional high frequency ultrasound (HFUS) imaging, as previously described (9,16). In brief, mice were anesthetized, the abdomen was depilated and B-mode image scans were acquired using a Vevo 2100 micro-ultrasound imaging system (Vevo LAZR; VisualSonics Inc., Toronto). Images were imported into Amira software (Visualization Sciences Group, Burlington, MA) for 3D tumor reconstruction and volume determination. Tumors developed in ∼100% of male animals for both GEMMs. The histology, lobular location and usual age to reach 300-500 mm^3^ volume are provided for the two models in Supplementary Table S1. Mice with tumor volumes of 300-500 mm^3^ were randomly enrolled into experimental groups, and surgically castrated (or sham-operated) via scrotal incision, under isoflurane anesthesia (17). To systematically define the response of the GEMMs to castration, we applied a murine approximation of the human clinical RECIST (response evaluation criteria in solid tumors) criteria (18). Thus, we define regression as a volume decline of 16% or more post-castration and recurrence (of tumors that had met the regression standard) as occurring when we observe volume increases of 11% from the volume nadir, in at least two temporally distinct imaging sessions. Etanercept (sTNFR2-Fc; Amgen, Thousand Oaks, CA, USA) was injected intra-peritoneally (4mg/kg) twice per week for up to 5 weeks following castration, based on our previous studies (8,11). The CCR2 antagonist (CCR2a) BMS-741672 (Torcis Bioscience, Bristol, UK) was injected intraperitoneally (2 mg/kg) three times per week as described (19), beginning one day prior to castration.

### Tumor tissue collection and processing

After mice were sacrificed, individual prostate tumors were identified grossly, excised and cut in half. Typically, a portion was flash frozen in liquid nitrogen and stored in a liquid nitrogen freezer while a second portion was fixed in paraformaldehyde at room temperature overnight, washed in PBS, stored at 4°C and embedded in paraffin as required for the preparation of tissue sections. Mouse prostate tumor and normal tissues were dissociated for single cell RNA sequencing using the Tumor Dissociation kits from Miltenyi Biotec (cat. no. 130-096-730). Briefly, tissue was excised, washed with chilled 1x PBS, placed in a petri dish on ice, cut into 2-4 mm^3^ pieces, transferred to a gentleMACS C tube (cat. no. 130-096-334) containing DMEM/enzyme mix and then attached to gentleMACS Octo Dissociator (cat. no. 130-096-427) with heating running the ‘37C_m_TDK_1’ (mouse) program. The suspension was filtered through a 40 µm cell strainer (Falcon). The cells were harvested by centrifugation at 400g for 10 min and resuspended in chilled 1X Red Blood Cell Removal Solution (Miltenyi Biotec, cat. no.130-094-183). Debris was then removed using the Debris Removal Solution (Miltenyi Biotec, cat. no. 130-109-398) following the manufacturer’s instructions.

### Cell culture

The C4-2 cell line, from Jer-Tsong Hsieh, UT Southwestern Medical Center, Dallas, TX, was used to model castration-resistant prostate cancer (20). C4-2 was periodically monitored by polymerase chain reaction (PCR) for mycoplasma contamination (21) and DNA fingerprinting was employed to authenticate this cell lines (22). C4-2 was cultured in RPMI-1640 media supplemented with 10% FCS and 1% penicillin/streptomycin. Enzalutamide was from Selleck Chemicals (Houston, TX).

### Enzyme-linked immunosorbent assays (ELISA)

In the case of prostate tumors, tissue powder was homogenized in NP-40 buffer using Sonic Dismembrator (Fisher Scientific, Hampton, NH) at power 7 for 30 sec. The supernatant was collected by centrifugation and stored at −80°C for subsequent analysis. In the case of cell culture, cells were seeded at a density of 0.8-1×10^5^/ml as mono-cultures and media stored at 4°C was used for subsequent analysis. TNF and CCL2 were quantitated by ELISA (eBioscience, San Diego, CA), according to the manufacturer’s instructions. Protein concentration was determined using bicinchoninic acid reagent (Pierce, ThermoFisher Scientific), according to the manufacturer’s instructions.

### Fluorescent activating cell sorting

Cells in culture were detached from cell culture plasticware using StemPro accutase (Gibco, Gaithersburg, MD), washed with PBS and incubated with 1% BSA in 1x PBS for 10 min at 4°C. Human cell lines were incubated with antibodies against CD166-FITC (3A6; Ancell, Stillwater, MN; 1:1250) and CD49f-Alexa Fluor^®^ 647 (GoH3; Biolegend, San Diego, CA; 1:200) for 30 min at 4°C. Mouse prostates were disaggregated using 1mg/ml type I collagenase and 1mg/ml Dispase (Invitrogen, Carlsbad, CA) in RPMI medium and filtered through 40 μm cell strainer. Cells were suspended in DMEM with 10% FBS and 1% L-glutamine, and then incubated with fluor-labeled antibodies for 30 min at 4°C. Mouse cells were incubated with antibodies against Sca-1-APC (D7; 1:500), CD49f-PE (GoH3; 1:333), and the ‘lineage’ markers (Ter119-FITC (TER-119; 1:250), CD31-FITC (390; 1:250) and CD45-FITC (30-F11; 1:250)). LSC FACS, first gating on lineage-negative (Lin^-^) cells and then CD49f and SCA-1 was performed as described by Witte and colleagues (23). All anti-mouse antibodies were purchased from eBioscience (San Diego, CA). 7-AAD (Invitrogen, Carlsbad, CA; 1:1000) was used to determine dead cells. FACS analysis was performed using the BD LSR-II (BD Bioscience, San Jose, CA) and analyzed by Winlist 3D (Version 8.0; Verity Software House, Topsham, ME, USA). Cell sorting was done by BD FACSAria-II (BD Bioscience, San Jose, CA).

### Colony formation assay

Cells were seeded in 6-well plates at densities of 0.5 × 10^3^, 1 × 10^3^ and 1.5 × 10^3^ cells per well and cultured for 21 days. The media was removed, wells were washed once with PBS, stained with 2% toluidine blue for 30 min and scanned with Epson Perfection 1240U (Suwa, Nagano Prefecture, Japan). Colonies were imaged and then enumerated using ImageJ.

### Immunohistochemistry

Immunohistochemical (IHC) staining was performed on 5 μm sections of formalin fixed paraffin-embedded (FFPE) mouse prostate tumor tissue. Sections were incubated with monoclonal antibodies against F4/80 (BM8; Biolegend, San Diego, CA, USA; 1:600), CD8alpha (D4W2Z; Cell Signaling Technology, Danvers, MA, USA; 1:400), or CCL2 (2H5; Biolegend, San Diego, CA, USA; 1:100), appropriate horse-radish peroxide secondary antibodies and reacted with 3,3-diaminobenzidine using an automated tissue processor. IHC and H&E images were captured at 20x magnification using Olympus BX45 microscope and cellSens software (Olympus, Shinjuku-ku, Tokyo, Japan). Ten random high powered (200X) fields were captured for each section and were blinded prior to quantitation of stained cells (F4/80^+^ or CCL2^+^ macrophages or CD8a^+^ lymphocytes) using ImageJ.

### RNA extraction, sequencing and analysis

RNA was extracted from liquid nitrogen frozen tumor samples by pulverizing tissue into a powder using a pre-cooled mortar and pestle. TRIzol (Invitrogen, Carlsbad, CA) was then added by a ratio of 1 mL of TRIzol per 100 mg of tumor tissue, and RNA extraction completed as per manufacturer’s instructions (24). Bulk RNA-seq data processing and analysis was performed as follows. FastQC v0.11.5 was used for quality control; STAR v2.6.0a (25) was employed to align the reads to the murine reference genome (GRCm38 M16) and gene expression was then quantified using RSEM (26). FastQC, RSEM, STAR were all performed in a local Linux/Unix computing environment. Subsequent analyses were performed using the following R-software packages from the Bioconductor website. Estimated gene counts from RSEM were filtered and the upper quartile was normalized using edgeR (27). Differential gene expression (DGE) was performed using Limma after Voom transformation followed by linear regression (28,29). The significance threshold was set to false discovery rate (FDR) / adjusted p-value of less than 0.05 with a fold change greater than 2 (FDR < 0.05 and |log2 fold change| > 1). Sets of genes with FDR<0.05 and |log2FC| from the DGE analysis were subjected to Gene Ontology analysis of biological processes as described (30), using the website tool (http://pantherdb.org/index.jsp). Heatmaps were generated using the package Complexheatmap (31). Single cell gene expression libraries were generated using the 10X Genomics Chromium platform and sequenced on a NovaSeq 6000 (Illumina). Processing of scRNA-seq data was performed as follows. For QC, normalization, clustering and differential gene expression, the Cellranger v6.0.0 software (https://support.10xgenomics.com/single-cell-geneexpression/software/) was used to align FASTQ sequencing reads to the mm10 (mouse) reference transcriptome, generating single cell feature counts for each sample. Count data from all cells were included in a ‘SingleCellExperiment’ (version 1.8.0) object (32). The ‘scater’ package (version 1.14.5) was used to filter cells and normalize counts per million (CPM) (33). Transcripts from a given cell were removed from the analysis if the corresponding transcriptome met any of the following criteria: 20% or more reads aligned to mitochondrial genes, 1000 total counts per cell or less, or 300 genes per cell or less. The dimensionality of this transcriptomic data was reduced using the t-distributed stochastic neighbor embedding (t-SNE) algorithm (34,35). The ‘limma’ package (29) was used to identify differentially expressed genes to refine the assignment of clusters generated by the t-SNE algorithm.

### Statistics

Data are presented as the mean ± SEM. Statistical analyses were conducted using JMP-Pro12 (SAS, Cary, NC, USA). Differences between two means was assessed by Student’s unpaired *t*-test and differences among multiple means by one-way or two-way ANOVA followed by Tukey-Kramer HSD post-hoc testing if the F-statistic was significant. For the small *n*, categorical data sets related to the studies using GEM prostate cancer models shown in Figures 4A, 5C and 5D, we employed the exact binomial test to determine the Clopper-Pearson confidence interval for the proportion and Fisher’s exact test to determine the *p*-values. Supplementary Table S3 summarizes these statistical analyses.

## RESULTS

### Tumor recurrence following castration-induced regression in PTEN-deficient PC

Mice engineered to delete the PTEN gene in the prostate epithelium at puberty uniformly develop an intraductal form of prostate adenocarcinoma (14) (Table S1). To investigate the response of prostate cancer to androgen deprivation therapies, we used high frequency ultrasound (HFUS) and three-dimension reconstruction software (9,16) to serially determine the volume of tumors in live animals (Fig. 1A). As seen in Figure 1B, the kinetics of tumor volume following castration is complex. Immediately following castration, tumor volume continues to increase similar to the non-castrated control. We recently demonstrated that this delay in tumor regression – which is not observed in the normal murine prostate – is driven by glucocorticoid receptor signaling transiently substituting for tumor-specific re-programmed AR signaling (9). Following this delay, the tumors regressed as expected but within ∼5 weeks most began to increase in volume, implying tumor recurrence and mimicking the development of CRPC in men. Regression and recurrence events were defined by adapting standard human oncology imaging protocols to mice, as described in the MATERIALS AND METHODS section and in the legend to Supplementary Table S2. Serial HFUS imaging of a large cohort castrated PTEN-deficient mice (including those in Fig. 1) is summarized in Supplementary Table S2, demonstrating 40-80% recurrence dependent on the period of post-castration observation.

**Figure 1.**
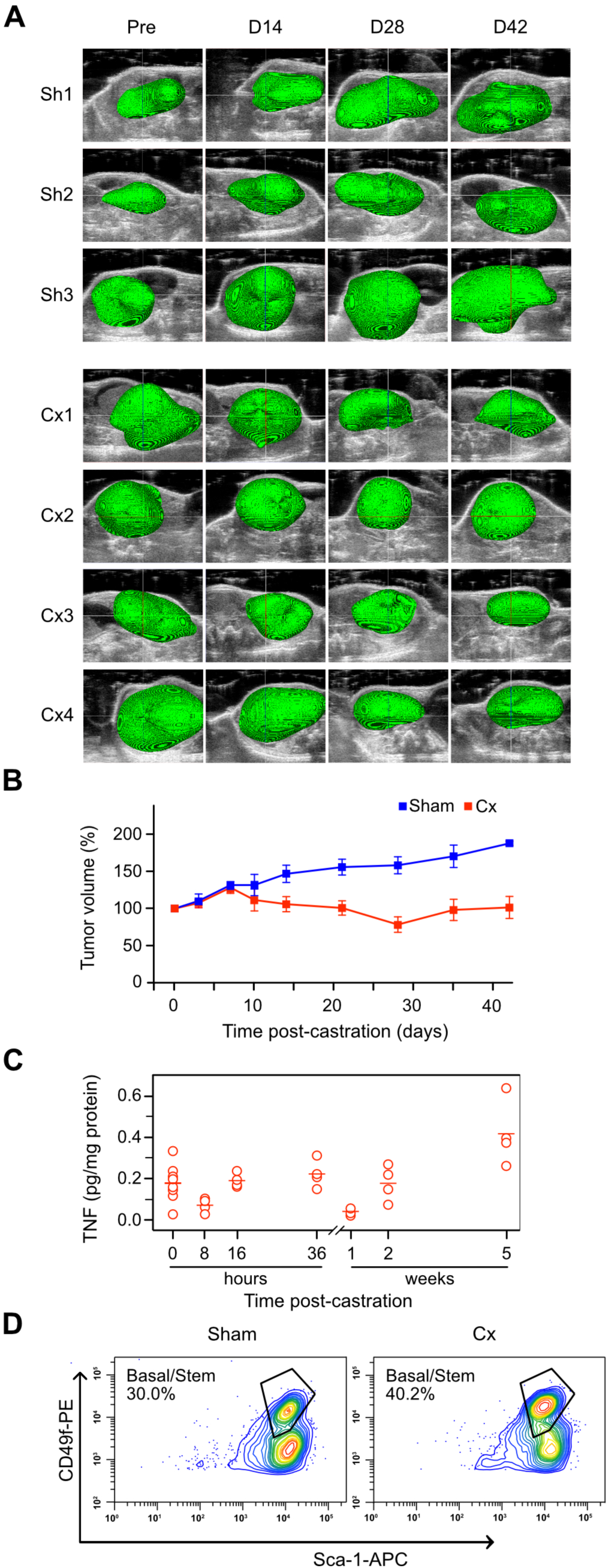
Castration induces tumor recurrence, and an increase in TNF and stemness in prostate tumors from PTEN-deficient mice. Mice bearing PTEN-deficient tumors were either castrated (Cx1-4) or sham-operated (Sh1-3) and serially imaged using HFUS. In some cases, tumor tissue was recovered for further analysis. **A**. 3D reconstruction of tumor-bearing ventral lobes (pseudo-colored green) performed as described in Materials & Methods. **B**. Tumor volume kinetics post-castration. Tumor volume was determined from HFUS images using Amira software. Mean tumor volume at the indicated times post-castration (n=4, red) or post-sham operation (n=3, blue) is shown as percent pre-castration volume (mean [%] ± SEM). **C**. TNF levels in tumors tissue extracts were measured by ELISA and normalized to total protein in the tissue sample, at the indicated times post castration. One-way ANOVA followed by Tukey-Kramer HSD test revealed a difference in TNF levels between 0 and 5 weeks (*p*<0.01). **D**. LSC FACS analysis of tumor cells from sham (left) and castrated (Cx; right) PTEN-deficient mice. Tumors were disaggregated and cells were processed for Lin^-^/Sca-1/CD49f FACS analysis (23). Relative fluorescence intensities are shown as contour plots. The LSC^hi^ population is labeled ‘Basal/stem’. The population in the lower right quadrant (LSC^med^) corresponds to luminal progenitors. A representative analysis is shown. Sham LSC FACS was replicated 5 times (mean[%]±SEM = 27.0±4.0); 25d post-castration LSC FACS was replicated 4 times (mean[%]±SEM = 38.2±1.7). Student’s t-test revealed a difference between Sham and Cx in the LSC^hi^ population (*p*<0.05).

We measured TNF protein levels in tumor lysates, and found an increase at 5 weeks post-castration, when tumor recurrence is beginning (Fig. 1C). Previously, we had observed an increase in *Tnf* mRNA at 8 hours post-castration in the stromal compartment of the normal prostate, suggesting a relatively direct effect of androgen-withdrawal on TNF expression (11). In the case of PTEN-deficient tumors, the increase in TNF protein at a late time post-castration suggests a more complex transcriptional and/or translational regulatory mechanism in response to androgen withdrawal. Alternately, the increase in TNF at late times post-castration might be due to the expansion of a population of TNF-expressing cells as the tumor evolves in response to androgen deprivation. We queried gene expression profiles in public databases from two previously published studies of men who had either hormone-naïve PC or CRPC (36,37), to gain insight into this question. Increased *TNF* mRNA expression in CRPC, relative to castration-naive primary PC, correlated with elevated expression of stem cell-related genes in both datasets (Fig. S1).

### Androgen-deprivation induced TNF secretion is due to increased basal cell stemness

To determine if we could detect a change in the stem cell-like tumor epithelial cell populations as the tumors began to increase in volume following castration-induced regression, we examined single cell preparations from tumors for cell surface expression of stem cell markers, using fluorescent-activated cell sorting (FACS). We observed an increase in Lin^-^/Sca1^+^/CD49f ^hi^ (LSC^hi^) cells at 25 days post-castration, relative to non-castrated tumors (Fig 1D). These cells correspond to the cytokeratin-5^+^/p63^+^ basal stem cell population (38), a multipotent stem cell population that is required for the post-natal development of the prostate (39). Basal stem cells are distinct from Lin^-^/Sca1^+^/CD49f ^med^ (LSC^med^) luminal progenitor cells (identified in the lower right population in Fig. 1D) that are the likely tumor initiating cell populations for most human prostate cancers (39,40).

The data in Figures 1 and S1 suggest that the TNF secreted during the post-castration recurrence phase in the PTEN-deficient tumors might be derived from a cell population with increased levels of ‘stemness’. To test this more thoroughly, we performed experiments using the C4-2 and LNCaP cell culture models of PC. First, we observed that long-term culture in the anti-androgen enzalutamide (41) significantly reduced the growth rate of C4-2 prostate cancer cells (Fig. 2A), as expected. Reduced growth was accompanied by an increase in the fraction of cells that resembled basal stem cells (Fig. 2B-C) and a coordinate increased in TNF levels (Fig. 2C). In the analyses for Figure 2, we employed CD166 as a basal stemness biomarker (42). We also performed the enzalutamide selection experiment on the LNCaP cell line, an isogenic parental line for C4-2 (20), which is more androgen-responsive (Fig S2A). We observed a similar correlation between stemness and TNF secretion (Fig. S2B-C). Finally, we replicated enzalutamide selection for both C4-2 and LNCaP cell lines, using immunofluorescence microscopy to identify CD49f^hi^ and CD166^hi^ cells (Fig. S3). Again, an increase in the fraction of cells scoring for these stemness markers correlated with increased TNF secretion. Stem cell marker enrichment >10% was typically required to detect an increase in TNF in the media of enzalutamide-treated cultures. Next, we treated C4-2 cells with enzalutamide for 20 days and used preparative FACS to partially purified cells into CD166^lo^ and CD166^hi^ fractions (Fig. 3A). TNF levels were higher in conditioned media from the CD166^hi^ fraction relative to the CD166^lo^ fraction (Fig. 3B). We also demonstrated that the efficiency of colony formation – a functional marker of stemness – correlated with the level of expression of CD166 (Fig. 3C).

**Figure 2.**
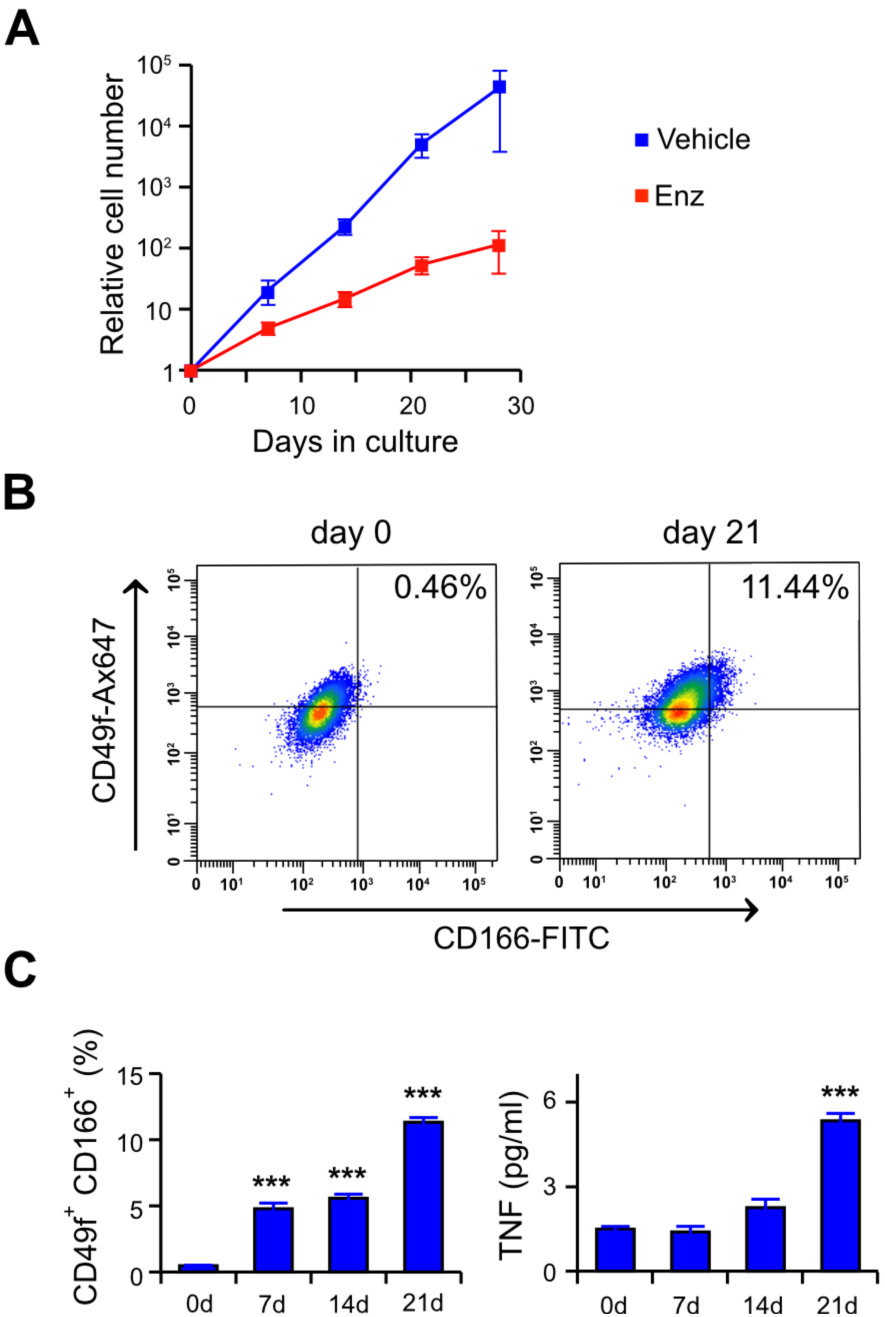
Extended enzalutamide treatment selects for basal stemness and TNF secretion. C4-2 cells were grown in media plus 10% serum and treated with vehicle (blue) or 10 μM enzalutamide (Enz; red) for the indicated time. **A.** C4-2 cell growth curve in the presence of enzalutamide. Trypan-blue excluding live cells were counted microscopically at the indicated times. Data are shown as mean ± s.e.m. (n=3). Two-way ANOVA followed by Tukey-Kramer HSD test revealed a difference in cell number for the Enz treated culture at 28d versus 0d (*p*<0.01). **B.** FACS analysis for basal cell stemness markers in enzalutamide treated C4-2 cells. Cells grown in enzalutamide for the indicated time, were incubated with the fluor-labeled antibodies, analyzed by FACS and relative fluorescence intensity represented as dot plots. The fraction of the cells that correspond to the CD49f^hi^/CD166^hi^ population (upper, right quadrant) is indicated. Additional time points are in Supplementary Figure S2. **C.** Fraction of cells scoring CD49f^hi^/CD166^hi^ and corresponding TNF secretion, at the indicated time of culture in media plus enzalutamide. Data from FACS analyses (see Fig. S2) is plotted on the left and TNF levels in the media determined by ELISA is plotted on the right. Values are given as mean ± s.e.m. (*n* = 2). **, *p* < 0.01 \*\*\**p* < 0.001 (two-way ANOVA followed by Tukey-Kramer HSD test).

**Figure 3.**
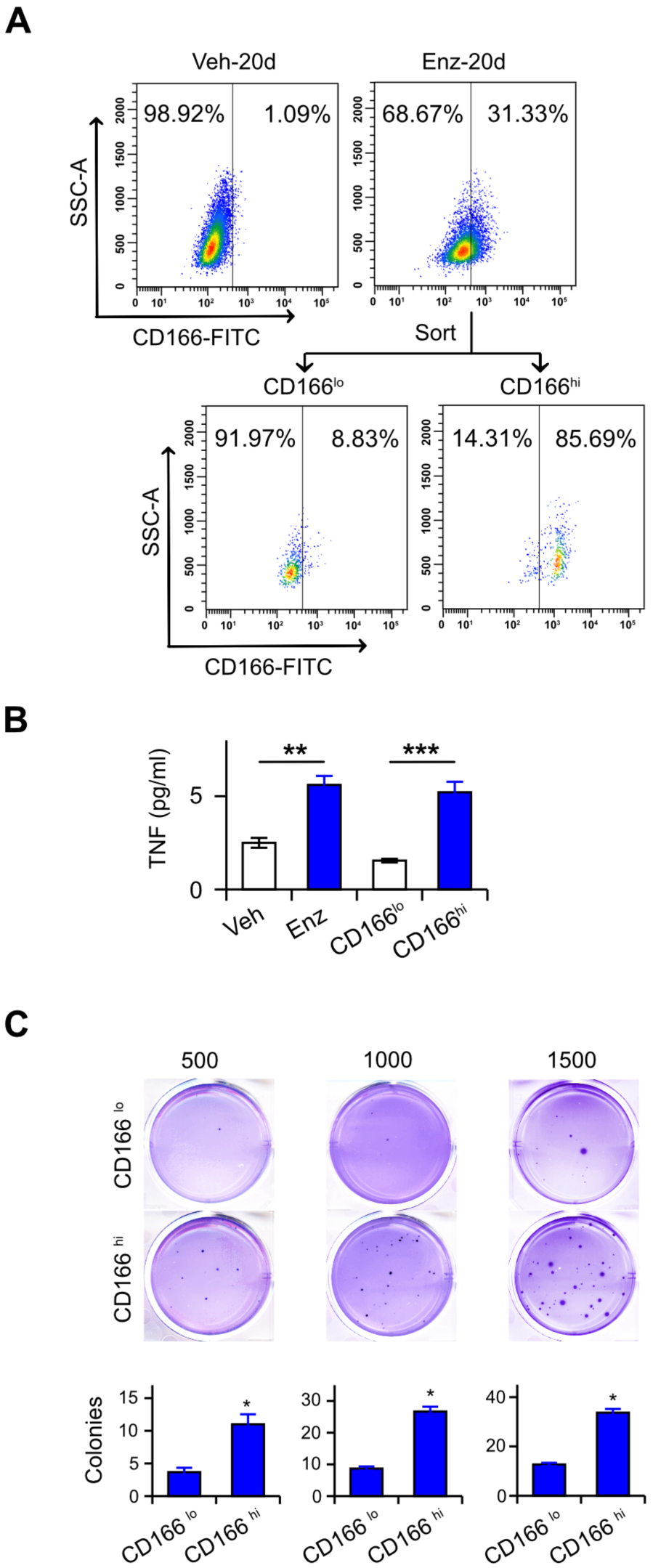
Enzalutamide treatment selects for a TNF-expressing basal stem cell population. C4-2 cells were grown for 20 days in media plus 10% serum containing 10 μM enzalutamide, enriched for CD166^hi^ expression. ELISA was used to measure TNF and colony formation. **A.** CD166-enrichment was performed via FACS to sort CD166^hi^ and CD166^lo^ C4-2 cells treated as indicated. The purity of the unsorted and sorted CD166^hi^ and CD166^lo^ populations are shown. **B.** TNF secretion of the indicated cultures (left to right): unsorted, vehicle-treated (veh); unsorted, enzalutamide-treated (enz); and sorted, enzalutamide-treated (CD166^lo^, CD166^hi^). TNF measured by ELISA, and plotted as mean ± s.e.m. (n=3). \*\**p*<0.01, \*\*\**p*<0.001 (One-way ANOVA followed by Tukey-Kramer HSD test). **C.** Colony formation assay of enriched CD166^hi^ and CD166^lo^ cell populations sorted as in **A**. Seeding densities were 500, 1000, and 1500 cells/well, respectively. Colony counts per well are plotted. \**p*<0.05 (Student’s unpaired *t*-test).

### Differentially-timed blockade of TNF signaling

We previously employed cell culture and mouse models of prostate cancer to demonstrate that TNF signaling mediates two distinct responses to ADT – an anti-tumorigenic apoptotic/regression phenotype (8,43) and a pro-tumorigenic cell migration phenotype (44). Given the observations in Figure 1, we hypothesized that TNF mediates both the regression and recurrence phases that occur after castration (Fig. 1B). To test this, we treated tumor-bearing mice with etanercept – a sTNFR2-Fc fusion protein, which competitively binds TNF (45) and prevents receptor binding – beginning at three distinct time points: three days prior, one day prior, and three days following castration. When etanercept is administered three days before castration (Fig. 4A, -3d), castration-induced regression is blocked, and tumor growth is similar to the sham group (Fig. 1B). Etanercept given one day before castration does not affect regression but blocks recurrence (Fig. 4A, -1d). Finally, when etanercept was administered three days after castration there is no detectable effect on regression or recurrence (Fig. 4A, +3d). Instead, the result resembles castration alone (Fig. 1B).

To gain insight into the nature of the different molecular processes occurring during regression and recurrence, we performed gene set enrichment analysis (GSEA) on RNA sequence data from bulk tumor mRNA. Specifically, we compared samples from sham castrated mice with those from either mice six days after castration (Fig. 4B, upper panel) or mice 35 days after castration (Fig. 4B, lower panel). Multiple gene sets related to TNF-induced transcription, most NFkB-related, were uniformly down-regulated at six days post-castration but up-regulated 5 weeks after castration, relative to the sham conditions. The switch in TNF-related gene sets from down-regulation to up-regulation is consistent with a switch from apoptosis signaling (low NFkB levels) to signaling that promotes transcription of NFkB-regulated genes (high NFkB levels).

**Figure 4.**
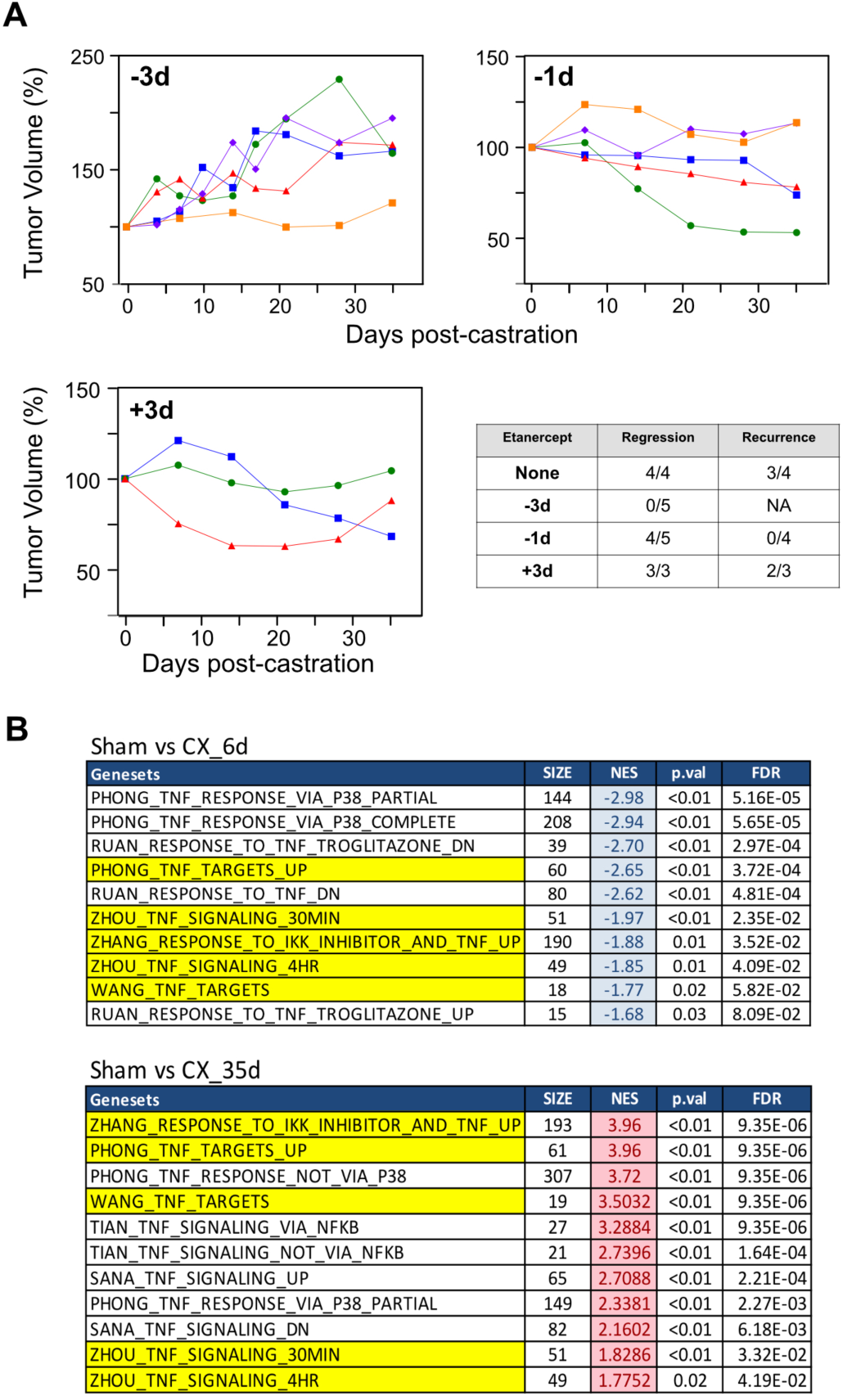
Differentially-timed TNF signaling blockade has distinct effects on tumor growth. Mice with PTEN-deficient tumors were castrated and etanercept (sTNFR2-Fc) was administered at the indicated times: three days before (-3d), one day before (-1d), or three days after (+3d) surgical castration. **A.** Tumor volume kinetics. Tumor volumes were determined by serial HFUS imaging and 3D reconstruction and plotted after normalizing to the initial tumor volume for each mouse. Different colors/symbols represent individual mice. The table summarizes regression and recurrence incidence for each experiment. Growth curves for castrated mice that did not receive etanercept (None) are not shown. The criteria for regression and recurrence are described in the Materials and Methods. Statistical analysis is provided in Supplementary Table S3A. **B.** Gene set enrichment analysis. RNA was extracted from PTEN-deficient tumors from the following mice: sham-operated, 6d post-castration and 35d post-castration. GSEA was performed on bulk RNA-seq data using gene sets corresponding to TNF-related pathways. Normalized enrichment scores (NES) from RNA from tumors at 6d (down-regulated, blue) and 35d (up-regulated, red) post-castration were compared to the sham group. Gene sets common to the two analyses are highlighted in yellow.

### CCL2 signaling is required for recurrence in murine prostate cancer models

CCL2 is a chemotactic cytokine (chemokine) (46) linked to prostate cancer pathogenesis (47). The CCL2-CCR2 signaling axis has multiple pro-tumorigenic functions, most notably the recruitment and polarization of tumor associated macrophages (TAMs) (48). Previously we demonstrated autocrine CCL2 secretion following enzalutamide induction of TNF in mono-cultures of C4-2 cells (10), consistent with CCL2 gene transcription via TNF induction of NFκB (49). We returned to the enzalutamide selection experiments shown in Figure 2C and Supplementary Figure S2 and found that the secretion of CCL2 protein increased over time, in a manner dependent on secreted TNF, in enzalutamide treated C4-2 and LNCaP cells (Fig. S4A-B)To assess paracrine CCL2 production in response to androgen deprivation, we replicated these experiments using co-cultures of LNCaP or C4-2 plus the myofibroblast-like WPMY-1 cell line (50), grown in enzalutamide-containing media. Again, we observed coordinate secretion of TNF and CCL2 (Fig. S4C). Co-cultures produced 5- to 20-fold more CCL2 than the prostate cancer cell line mono-cultures. We also found that enzalutamide-treated co-cultures of C4-2 plus WPMY-1 produced conditioned media that induced migration of THP-1, a CCR2^+^ macrophage-like leukemia cell line that is a model for TAMs (Fig. S4D).

These observations suggested that paracrine TNF induction of CCL2 in the prostate cancer TME might be responsible for tumor recurrence at late times post-castration, possibly by recruiting immunosuppressive TAMs (51). Indeed, we found that in the samples from Figure 1C, CCL2 increased at 5 weeks (Fig. 5A), coincident with the rise in TNF that accompanies tumor recurrence. Tissue sections corresponding to tumors 5 weeks post-castration showed an increase in stroma-localized macrophage-like cells expressing CCL2 (Fig. 5B). Moreover, CCL2 expression was reduced in tumor tissue sections from castrated mice that received etanercept, consistent with CCL2 regulation by TNF (Fig. 5B). To determine if CCL2 signaling was necessary for post-castration tumor recurrence in prostate cancer mouse models, mice bearing PTEN-deficient prostate cancers received an antagonist of CCR2 prior to castration. Serial imaging revealed that none of the tumors met the criteria for recurrence (Fig. 5C), indicating that CCL2-CCR2 signaling is indeed required for castration resistant recurrence in this model. Importantly, the tumor kinetics produced by CCR2 blockade phenocopies the effect of etanercept administered one-day prior to castration (Fig. 4A, -1d), consistent with TNF regulation of CCL2 gene transcription via NFkB. To confirm the role of CCL2 signaling we replicated the CCR2 inhibition studies in the Hi-MYC prostate cancer GEMM, which has a similar response to castration (regression delay, regression then recurrence) as the PTEN-deficient GEMM (Fig. 5D, blue symbols; see also ref. (9)). We observed that CCR2a treatment prior to castration was comparably effective at preventing post-castration recurrence in Hi-MYC mice (Fig. 5D, red symbols; also see Fig. S5).

**Figure 5.**
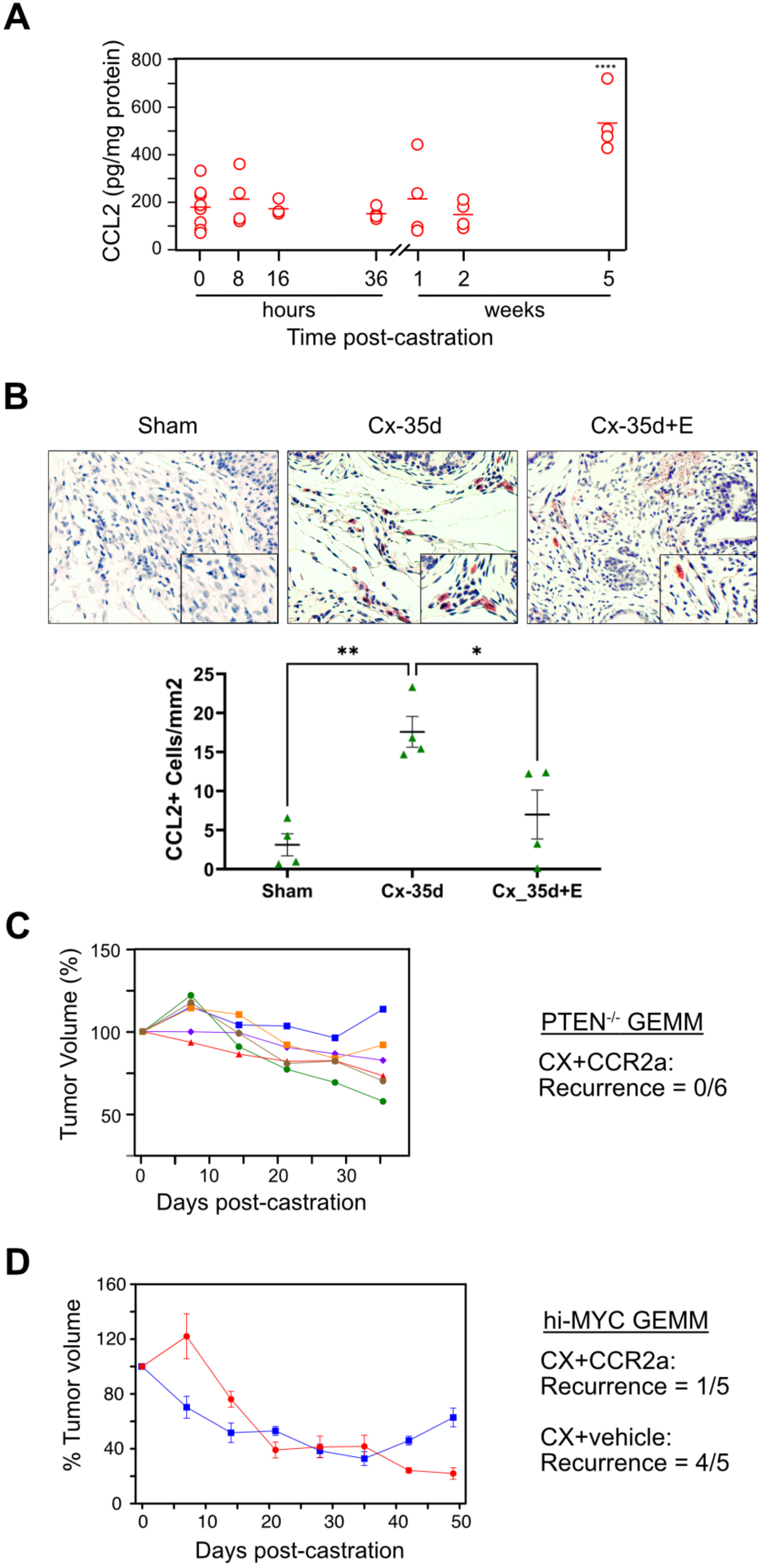
CCL2-CCR2 signaling is required for castration-resistant recurrence. Mice bearing PTEN-deficient (**A-C**) or hi-MYC (**D**) tumors were sham-operated or castrated at the indicated times. Volume was monitored by HFUS. Some mice received etanercept (E) or the CCR2a, as indicated. **A**. Tumor CCL2 levels following castration. CCL2 levels were measured by ELISA on the same tumor extracts prepared for Fig. 1C, normalized to total protein and plotted as open circles. The bar marks the mean. Two-way ANOVA followed by Tukey-Kramer HSD test revealed a difference in CCL2 levels between 0 and 5 weeks *(p* < 0.0001).( **B.** Representative images of IHC-stained CCL2^+^ cells (red) in tumors from the indicated treatment groups. CCL2+ cells were counted and plotted. \*\**p* < 0.01, \**p* < 0.05 (Two-way ANOVA followed by Tukey-Kramer HSD test). **C.** Treatment with a CCR2 antagonist phenocopies -1d etanercept treatment (Fig. 4A) in PTEN-deficient tumor-bearing mice. Tumor volumes were determined by serial HFUS imaging and 3D reconstruction and plotted after normalizing to the initial tumor volume for each mouse (left). Different colors/symbols represent individual mice. The recurrence incidence is summarized (right). **D.** CCR2 antagonist suppresses castration-induced regression in Hi-MYC prostate cancer-bearing mice. Blue boxes: mice were castrated and received vehicle. Red circles: mice were castrated and received CCR2a prior to castration. N=5 for each group. The recurrence incidence is summarized (right). Tumor volume kinetics for individual mice are in Supplementary Figure S5. Statistical analysis for **C-D** is provided in Supplementary Table S3B.

We used a transcriptomic approach to further investigate CCL2 signaling, immune cell migration, and TAMs in the development of castration-resistant prostate cancer in the PTEN-deficient GEMM. Differential gene expression (DGE) analysis was performed on bulk RNA-sequencing data from tumor tissue and genes with FDR<0.05 and |log2FC| (see Fig. S6 and details in the legend) were subjected to Gene Ontology (GO) enrichment analysis (Fig. 6). These analyses revealed that a very similar category of gene sets – mediating immune cell chemotaxis and migration – are up-regulated in the bulk tumor mRNA from mice 35 days post-castration (Fig. 6A) but are down-regulated in the bulk tumor mRNA from mice 35 days post-castration which were treated with CCR2a (Fig. 6B). We also assembled a TAM-related gene set and found that these genes are similarly regulated: ‘up’ in the 35 days post-castration group and ‘down’ in the 35 days post-castrated group treated with CCR2a (Fig. 6C).

**Figure 6.**
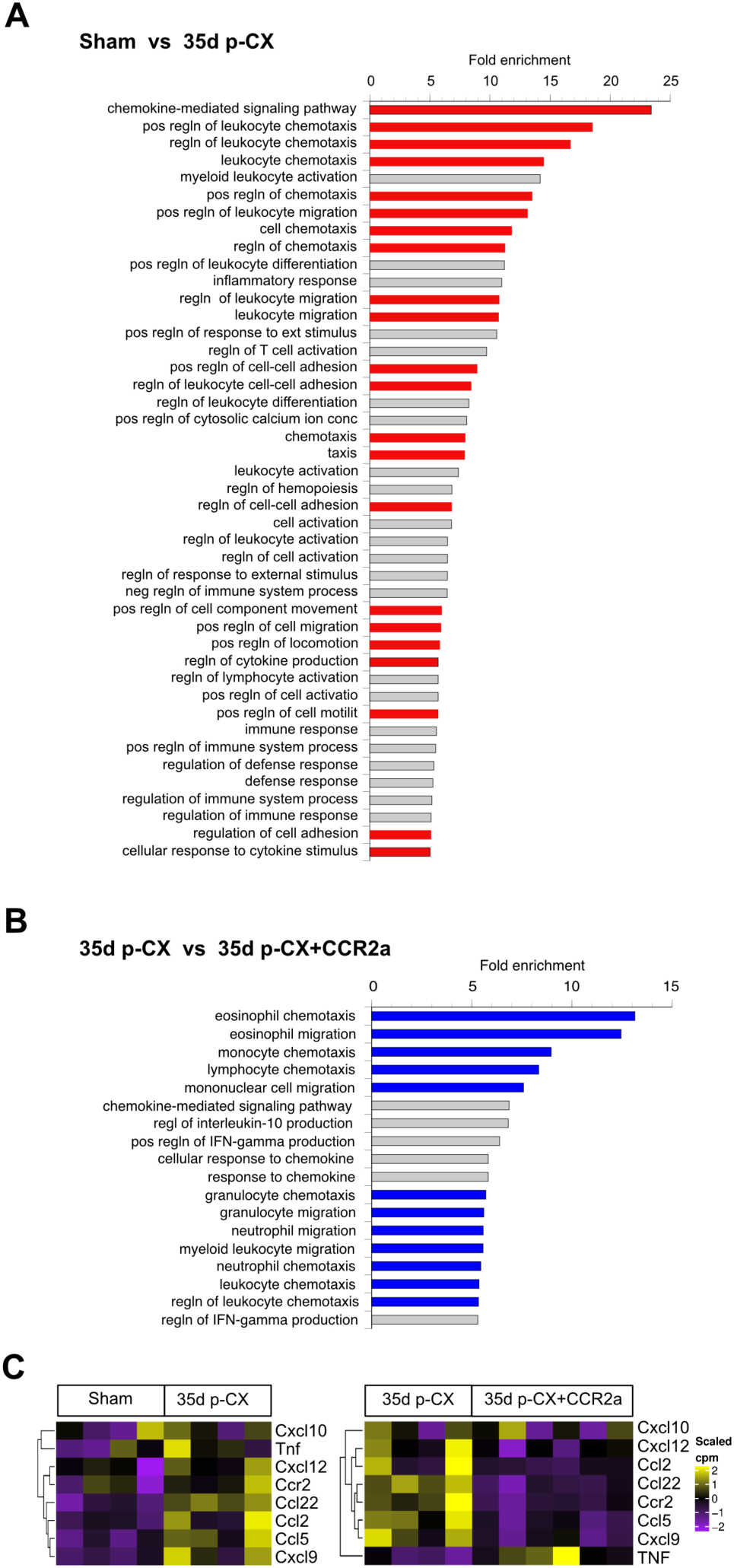
Transcriptomic changes in PTEN-deficient prostate cancers following CCL2 signaling blockade. RNA was extracted from PTEN-deficient tumors from the following mice: sham-operated (Sham), 35d post-castration (35d post-CX) and 35d post-castration after receiving CCR2a (35dp-CX+CCR2a). DGE analysis was performed on bulk RNAseq data (Fig. S6) followed by Gene Ontology analysis of biological processes comparing RNAseq transcript sets derived from tumors in mice that were sham-operated vs mice 35d post-castration (**A**) or from tumors in mice 35d post-castration vs mice treated with CCR2a 35d post-castration (**B**). Red-colored bars represent up-regulation and blue-colored bars represent down-regulation. Fold-enrichment >5. **C.** TAM-related transcripts regulated by CCL2 signaling. Heatmap representation of mRNA levels of select TAM-associated genes for the same two comparisons shown in **A** (left) and **B** (right).

### An immunosuppressive cell profile is induced in the TME during recurrence

Taken together, the analyses in Figures 5 and 6 indicate that CCL2 might be functioning to recruit TAMs and drive the progression to, or recurrence as, castration resistant prostate cancer (52). Thus, to determine how TNF-CCL2 signaling effects the tumor immune microenvironment in the PTEN-deficient murine prostate cancer model, we used immunohistochemistry (IHC) to quantitate F4/80- and CD8-stained cells in tissue sections from the tumors. As seen in Figure 7A, F4/80-stained cells (presumptive TAMs) increase, most notably at 35 days post-castration, just as tumor re-growth in the recurrence phase is beginning and CCL2 levels are peaking (Fig. 5A). In addition, etanercept partially blocks this increase in TAMs and coordinately inhibits tumor recurrence. CD8 T cells (presumptive cytolytic T-lymphocytes; CTLs) decreased over time following castration and this was also partially reversed by etanercept administration to the mice bearing PTEN-deficient tumors. Figure 7B demonstrates that blocking CCL2 signaling with the CCR2 antagonist has the expected effect – it reduces F4/80-stained cells and increased CD8-stained cells, reversing the castration-induced immunosuppressive cell profile and coordinately preventing prostate cancer recurrence (Fig. 5C). Integrating the data on F4/80-staining, CD8-staining and CCL2 ELISAs for tumors treated and harvested at various times post-castration reveals a demonstrates a high degree of correlation of the key immune components we examined. Specifically, there is positive correlation of macrophages and CCL2 secretion and a negative correlation of T cells with macrophage as well as with CCL2 section (Fig. S7). We also observed a more pronounced CD45+ cell infiltrate in the recurrent tumors, relative to non-recurrent tumors (Fig. S7B).

**Figure 7.**
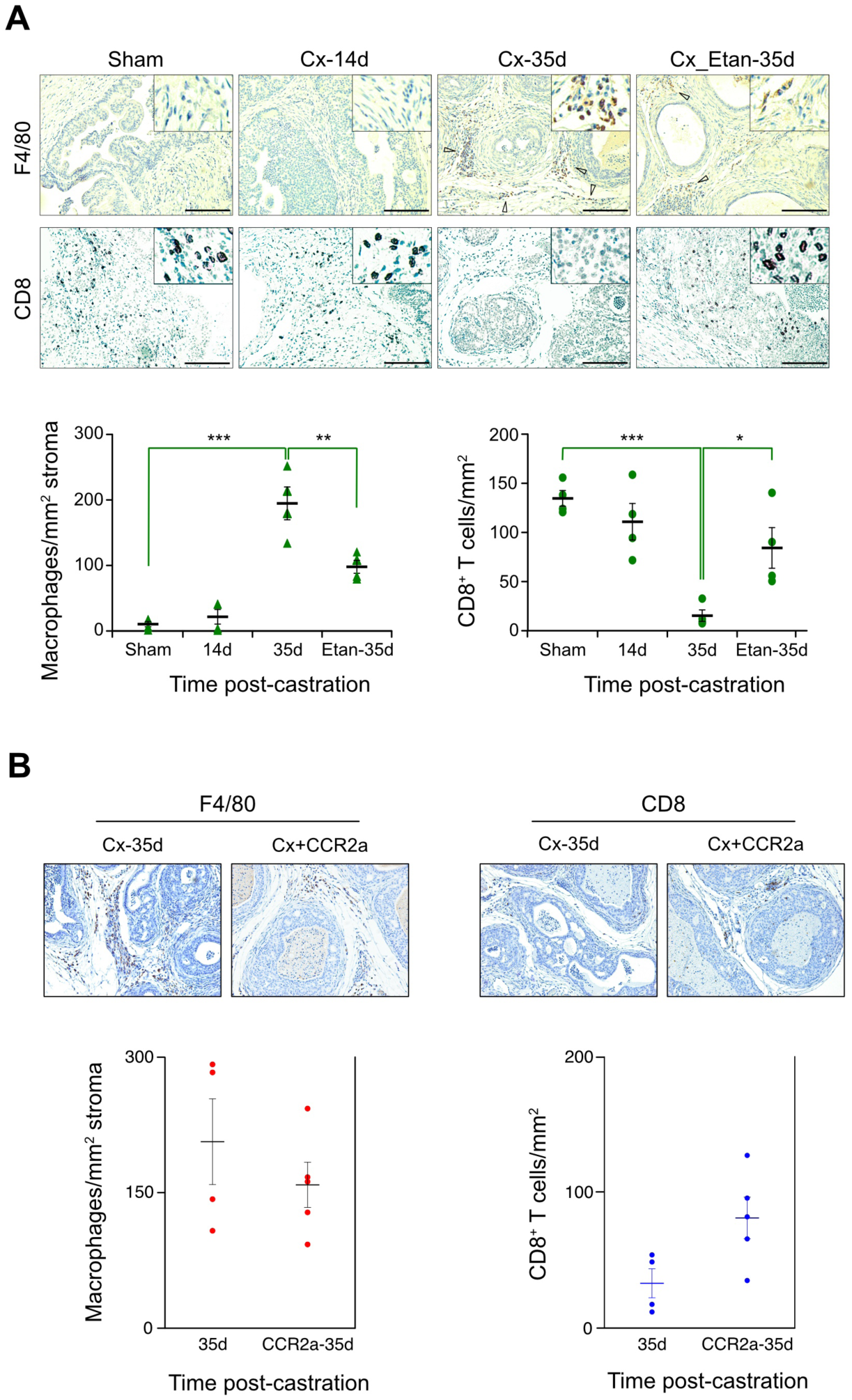
TNF and CCR2 signaling blockade suppress castration-induced immunosuppression at late times following castration. Mice bearing PTEN-deficient tumors were sham-operator or castrated (Cx) and tumors harvested at the indicated times post-castration. Some mice received etanercept (Etan) or the CCR antagonist BMS IHC staining for stromal F4/80+ macrophages and CD8+ T cells was performed and the number of stained cells per unit area were plotted. **A**. TNF signaling blockade. Images of representative IHC-stained tissue sections for the surface marker protein noted on the left side. Mice were treated as shown across top and the cell densities plotted for each of the treatments, below. **B**. CCL2 signaling blockade. Similar to **A** with both surface marker protein and corresponding mouse treatments shown along the top. A complete set of images are provided in Supplementary Figure S7C. Staining and counting of the Cx-35d samples in **A** and **B** were performed separately. Data are given as mean ± s.e.m. (n=4). *p<0.05, **p<0.01, ***p<0.001 (One-way ANOVA followed by Tukey-Kramer HSD test). *P*-value for the differences for CD8+ T cells and F4/80 macrophages due to CCR2a treatment was >0.05 (not statistically significant).

### Transcriptomic CD8/CD68 ratio predicts castration outcome

Next, we carried out transcriptomic profiling to confirm the IHC studies in Figure 7, which correlated an immunosuppressive immune cell profile (elevated CD68, reduced CD8) with recurrence of the PTEN-deficient tumor. Specifically, three mice bearing PTEN-deficient prostate cancers were castrated and tumor volume was HFUS-monitored for 42 days (Fig. 8A). One mouse failed to regress and was labeled ‘stable’. Two mice regressed, one of which showed recurrence 21 days after castration. We recovered the tumors, and performed single cell RNA sequencing. Transcript count data was pooled and tSNE dimensionality reduction was used, along with subjective cell annotation, to identify six cell populations (Fig. 8B). Among CD45-positive immune cells, TAMs were identified as those cells expressing both CD68 and CD80, while cytolytic T cells (CTL) were defined as those expressing CD3 and CD8 (see details in the legend to Fig. 8). In Figure 8C, we plotted the ratio of CTL to TAM, and observed that compared with the regressing tumor, the recurrent tumor had a lower CTL/TAM ratio. This is consistent with the IHC analysis in Figure 7A and suggests that immune suppression drives tumor re-growth.

**Figure 8.**
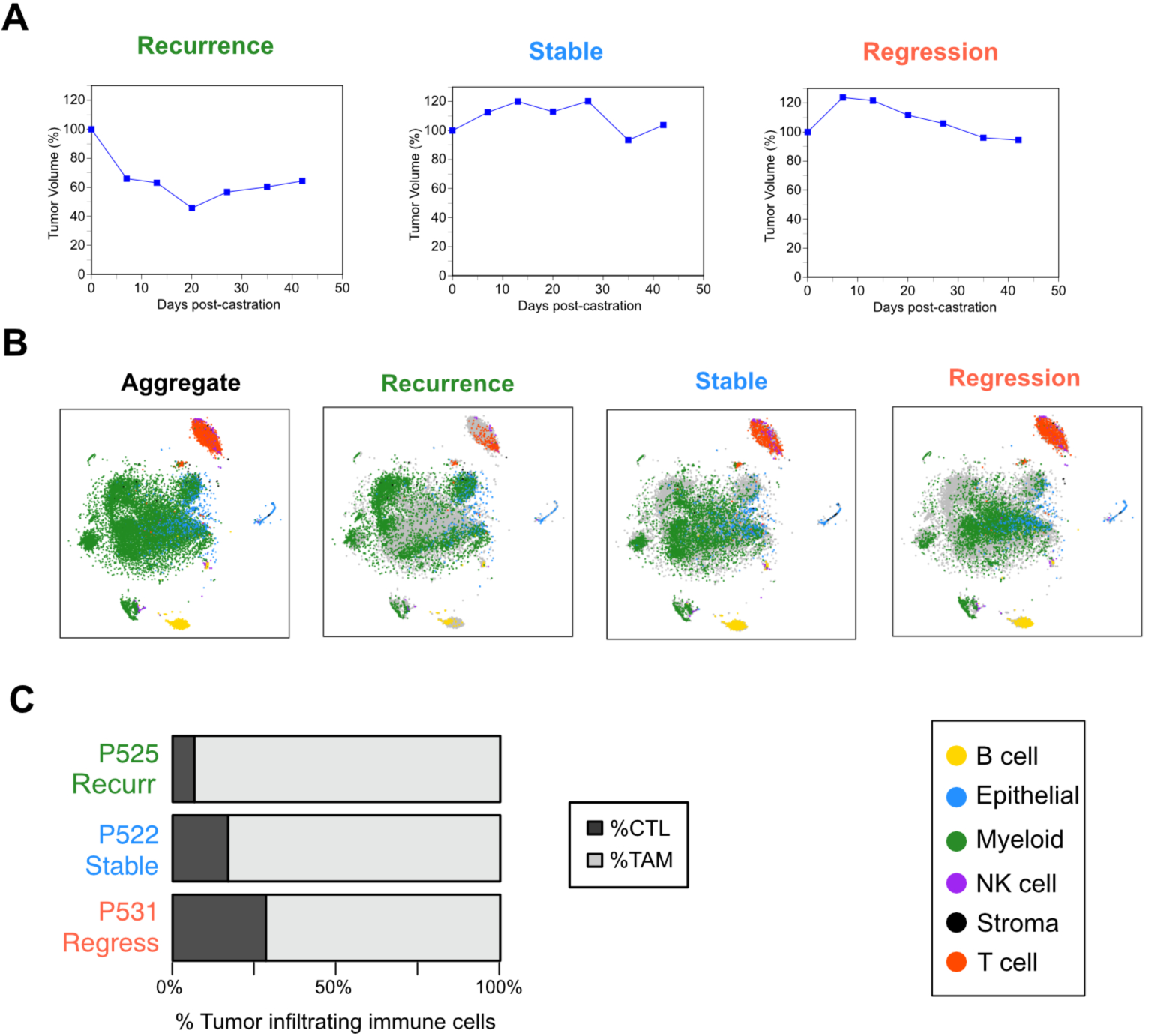
Tumor recurrence following castration correlates with an immunosuppressive tumor microenvironment. **A.** Post-castration tumor volume kinetics for three representative PTEN-deficient mouse tumors. Tumor volumes were determined by serial HFUS imaging and 3D reconstruction and plotted after normalizing to the initial tumor volume for each mouse. The average starting volume for these tumors was ∼960 mm^3^. **B.** t-SNE plots for single cell RNAseq analysis. At 42d post-castration, mice were sacrificed, tumor tissue was disaggregated into single cell preparations, RNA sequenced and analyzed as described in the Methods. Dimensionality reduction of cell transcriptomes from all three mice was performed with tSNE to produce the plot in far left panel (Aggregate). Cells are assigned to 6 subset populations as coded in the key in the lower right corner. Color-coded cells from each of the three mice corresponding to the response patterns in **A** are shown in the remaining three panels of **B**. Cells which were in the aggregate population, but not identified in individual mice are grey colored. TAMs (defined in panel C) are within the large, heterogeneous myeloid population (green). **C.** CTL:TAM ratios for regressing, stable or recurrent mouse tumors. Transcriptomic data from **B** was used to define TAMs (CD45 and CD68and CD80; lighter grey) and CTLs (CD45 and [CD3e or Cd3d or CD3g] and [CD8a or CD8b1]; darker grey). See text for discussion.

## DISCUSSION

### TNF signaling switch

Employing a HFUS-based imaging protocol to measure tumor volume in live mice, we are able to resolve the response of genetically-engineered murine prostate cancers to castration into multiple, mechanistically-distinct phases, as illustrated for the PTEN-deficient prostate cancer model in Figure 1 (the kinetics for the castration response in the Hi-MYC GEMM are similar). Immediately following castration, there is an increase in tumor volume in both prostate cancer GEMMs. We recently demonstrated that this seemingly paradoxical effect is dependent on glucocorticoid signaling transiently substituting for tumor-specific androgen receptor signals that are blocked by the androgen ablative effects of castration (9). After 4-7 days of gradually increasing volume, the tumors begin to regress due to apoptotic cell death of the epithelial tumor cells in the cancerous glands, consistent with the classical morphological description of apoptosis (53). In addition to tumor epithelial cell apoptosis, there is a loss of vascular integrity within the tumor that contributes to the regression phase (54,55). Some tumors continue to regress for more than two months, but for the majority – up to 75% of individual tumors in the prostate cancer GEMMs we studied – recurrence begins 4-6 weeks after castration (Fig. 1A, Fig 4D and Supplementary Table S2b).

We sought to decipher the intra- and inter-cellular signaling events that occur within the tumor and the surrounding microenvironment in response to androgen deprivation (castration). Previously we had shown that TNF – but not other common death receptor ligands (FasL and TRAIL) – is necessary for apoptosis-driven regression of the normal murine prostate (11). In addition, we also found that TNF is required for so-called vascular regression of murine prostate cancers, probably by triggering endothelial cell apoptosis (8). The loss of vascular integrity leads to tumor hypoxia within a day following castration. While these events occur multiple days prior to the onset of a measurable reduction in prostate cancer volume, hypoxia may indirectly drive regression by inducing tumor epithelial cell apoptosis via TP53 (8). In the current studies, when we administered etanercept to PTEN-deficient mice three days prior to castration, regression was effectively inhibited (Fig. 4A and Supplementary Table S2a). This observation is consistent with TNF-driven apoptosis of tumor epithelial and endothelial cells.

The increase in TNF protein 5 weeks post-castration (Fig. 1C) – at a point when most tumors have regressed – suggested a second role for TNF signaling in the response to androgen deprivation. Indeed, blockade of TNF signaling beginning one day prior to castration results in most tumors regressing, but no recurrence (Fig. 4A and Supplementary Table S2a). Thus, TNF signaling switches from pro-apoptotic signaling that activates caspase-8 and drives regression to pro-tumorigenic signaling that activates the NFκB and/or PI3K/Akt pathways, promoting tumor cell survival and proliferation and tumor recurrence. The results of transcriptional analysis (Fig. 4B) support the switch to a pro-tumorigenic phenotype: we observed increased expression of NFκB-related gene sets in bulk mRNA recovered from tumor samples at 35 days, versus 6 days, post-castration. A key consequence of NFκB signaling during the recurrence phase is the induction of CCL2 protein levels 35 days post-castration (Fig. 5A). Moreover, the administration of a CCR2 antagonist to tumor bearing mice prevented castration-induced recurrence (Fig. 5C), phenocopying the effects of -1d TNF blockade by etanercept treatment (Fig. 4A, -1d panel). This observation is consistent with CCL2 expression downstream of TNF-NFκB signaling. Similarly, the number of CCL2-staining cells in the TME was reduced in tumors from mice receiving etanercept to block TNF signaling (Fig. 5B).

### Increased tumor cell stemness following androgen deprivation drives pro-tumor TNF signaling

While the pleotropic signaling effects of TNF are well-established, an important mechanistic question in the context of this report is whether changes in the tumor and surrounding TME are responsible for the post-castration switch in TNF signaling. Two potential determinants are the spatial and temporal pattern of TNF production/ secretion and the context of the cells that TNF signaling targets. Previously, we showed that in the normal rat prostate TNF mRNA and protein increased as early as 8 hours post-castration, with the most significant increase occurring within the stromal compartment of the prostate (11). However, in PTEN-deficient murine tumors there is no discernable increase in TNF protein in the first 24 hours post-castration (Fig. 1B). Alternatively, death receptor driven apoptosis can be triggered by down-regulation of the CASP8 inhibitor CFLAR (56). Indeed, we previously found castration reduced CFLAR expression within 12 hours post-castration in the normal rat prostate (12,13). Moreover, CFLAR protein is up-regulated in CRPC patient samples (57). To investigate the mechanism of the late (5 weeks post-castration) increase in TNF, we blocked AR action with enzalutamide in prostate cancer cell culture models (10). After 3 weeks of culture in enzalutamide, TNF secretion and stemness were increased in both C42 and LNCaP cell lines (Figs. 2 and Supplementary Figs. S2-S3). Higher levels of TNF secretion occurred in cultures enriched for expression of one of the stem cell-like markers (Fig. 3). The increased expression of stem cell-like genes following androgen deprivation may reflect the death of luminal-like cells (low stemness) and selection for, or expansion of, basal-like cells (higher stemness) that have increased levels of NFκB and therefore secrete more TNF protein. We observed an increase in basal stem-like cells at 25 days post-castration in the PTEN-deficient tumor model (Fig. 1D), suggesting a similar phenomenon is occurring *in vivo*. It is also possible that the increase in stemness that we observe post-castration is due to lineage plasticity of pre-existing cells, rather than selection and expansion of *bone fide* tumor stem cells (58). Regardless, it appears most likely that the increase in TNF secretion observed 5 weeks post-castration is a consequence of changes in cellular context rather than a promoter/enhancer-driven transcriptional response.

### Immune suppression is a key consequence of increased tumor cell stemness

By investigating the switch in TNF signaling, we identified CCL2 secretion as a key TNF-NFκB-induced event required for the onset of tumor recurrence in the PTEN-deficient GEMM (Fig. 5). Moreover, CCL2 is also required for recurrence in the genetically distinct Hi-MYC GEMM, suggesting CCL2 may be a common driver for recurrence post-castration, at least in murine models of prostate cancer. Although CCL2 has multiple pro-tumorigenic immune-modulating functions in prostate and other cancers (59), the transcriptomic studies described in Figure 6 are consistent with TAM recruitment. Specifically, Gene Ontology analysis (Fig. 6A-B) indicates that CCL2 induces immune cell chemotaxis and migration at the time of recurrence, which is the behavior expected of TAMs as well as other myeloid immune suppressive cells, such as myeloid-derived suppressor cells (MDSC). The heat map in Figure 6C showed up-regulation of TAM-related genes, while IHC analysis in Figure 7 (and Supplementary Fig. S7) demonstrated a coordinate increase in F4/80+ macrophages and CCL2 at the time of recurrence. The latter two observations support a critical role for TAMs, but do not rule out a role for MDSC. Identifying the precise tumor immune cell populations within the TME that modulate the response of cancers to therapy is the focus of considerable ongoing research (60). TAMs are a heterogenous set of immune cells which are predominantly pro-tumorigenic (61) and have been implicated in human and murine prostate cancers (62,63). Although an oversimplification of the immune response to cancer, a balance between anti-tumor cytolytic T cells and immunosuppressive myeloid cell populations may predict the success of emerging immunotherapies. Indeed, variants of the CTL/TAM ratio – similar to our observations in Figures 7 and 8 – are potential prognostic biomarkers in patients with melanoma (64), breast cancer (65) and hepatocellular carcinoma (66).

### Prospective: a model for the development of CRPC

The models we employed are anatomically similar to non-metastatic CRPC (nmCRPC) in that the tumor is localized to the prostate, whereas in metastatic CRPC (mCRPC) the TME is derived from the metastatic site, such as bone. Most CRPC is discovered clinically as mCRPC and indeed nmCRPC is likely a transient state that progresses to mCRPC (67). Castration of mice bearing PTEN-deficient or Hi-MYC prostate cancers induces regression and then recurrence – a pattern very similar to that seen in the development of human CRPC. Importantly, HFUS imaging provides real-time volume data to ensure that the complex kinetics of tumor volume changes can be used to accurately categorize the response of each tumor to androgen deprivation. Supplementary Figure S8 is a mechanistic model consistent with our findings. It proposes that TNF secretion into the TME first drives regression and subsequently induces an immunosuppressive state that leads to recurrence. The specific signal may vary depending on the TME context. For example, in some PTEN-deficient GEMMs, IL1ß triggers myeloid immune suppression (68). Most of our analyses were performed on bulk tumor tissue – this represents a significant limitation of the studies in this report. Future studies employing multi-modal single cell analyses will be critical to elucidating the molecular interactions between tumor cells and the myofibroblasts, endothelial cells and immune cells that constitute the TME. These studies are underway in our laboratories. Although the CCL2 antagonist was effective at preventing recurrence in our GEMMs, a monoclonal antibody which binds CCL2 failed in a phase 2 patient trial in mCRPC (69). This may have been due to technical issues with the drug (69), or possibly heterogeneity among human CRPCs. Indeed, Giancotti and colleagues identified three subtypes of CRPC (70) and proposed that one of these – mesenchymal stem-like prostate cancer – might use CCL2 signaling to promote myeloid immune suppression. Thus, in the future patients with this CRPC subtype may be candidates for anti-CCL2 therapies.

## ACKNOWLEDGEMENTS

We acknowledge: Chawnshang Chang and Shu-Yuan Yeh (University of Rochester) for encouraging us to investigate the pro-tumorigenic role of CCL2 in prostate cancer; Yvonne Saenger (Albert Einstein College of Medicine) for helpful discussions on the ratio of CTLs and TAMs in tumor immune suppression, Bo Xu (Roswell Park Comprehensive Cancer Center) for discussions on the histopathology of murine prostate cancers and Michalis Mastri (Thermo-Fisher/ GIBCO) for assistance with computational analysis of RNA sequence data.

## Notes

**FINANCIAL SUPPORT:** This work was supported by the Department of Defense Prostate Cancer Research Program No. W81XWH-19-1-0378 (to J.J. Krolewski and K.L. Nastiuk), institutional funds from the Roswell Park Alliance Foundation and the National Cancer Institute Cancer Center Support Grant P30-CA016056 to Roswell Park Comprehensive Cancer Center.

### Competing Interest Statement

The authors have declared no competing interest.

### Summary of Updates

Addition of supplementary data to Supplementary Figure S4 (Panel B); addition of statistical analyses including Supplementary Table S3; minor modifactions and corrections to the text.

