## Supplementary Data for "Androgen deprivation triggers a cytokine signaling switch to induce immune suppression and prostate cancer recurrence"

### SUPPLEMENTAL METHODS

#### Cell culture

LNCaP, WPMY-1 and THP-1 cell lines were from the American Type Culture Collection (Rockwell, MD, USA). Cells were grown in RPMI-1640 media supplemented with 10% fetal calf serum and 1% penicillin/streptomycin. For THP-1, the media was further supplemented with 10 mM HEPES and 2 mM glutamine. Cell lines were periodically monitored by polymerase chain reaction (PCR) for mycoplasma contamination (21) and DNA fingerprinting was employed to authenticate cell lines (22). LNCaP is an androgen sensitive cell line that is used to model primary prostate cancer (71). WPMY-1 is an SV40 T-antigen immortalized cell line derived from a human cancer (50). It expresses smooth muscle actin and vimentin, and is a model for prostate stromal myofibroblasts. THP-1 is derived from a human acute monocytic leukemia (72). It is CCR2-positive (73), displays monocytic cell surface markers and has phagocytic activity, making it a model for tumor associated macrophages (74).

#### Analysis of public DNA microarray gene expression datasets

The web browser interface of the Gene Expression Omnibus (GEO) database (www.ncbi.nlm.nih.gov/geo/) or the ONCOMINE gene expression database (www.oncomine.org), were used to extract relevant DNA micro-array expression datasets and compare relative gene expression levels for primary and castration-resistant prostate cancers. The results were visualized as box and whiskers plots.

#### Immunofluorescence microscopy

C4-2 and LNCaP cells were seeded onto cover slips (Corning Inc., Corning, NY, USA; 18mm x18mm) in 3.5cm tissue culture plates at a density of 20,000 cells/slide. Cells were cultured overnight to allow adherence to the coverslips, and then fixed in ice-cold methanol for 10 minutes. Cells were permeabilized with 0.1% Triton-X-100 (Sigma-Aldrich, St. Louis, MO, USA) in 1x PBS for 5 minutes, washed with PBS and incubated for 1 hour with 5% BSA in 1x PBS to block non-specific binding. Cells were then incubated with CD49f-Alexa Fluor® 647 antibody (GoH3; Biolegend, San Diego, CA, USA; 1:100) overnight at 4°C, or stained with a primary anti-CD166 antibody (MOG/07; Leica Biosystems, Wetzlar, Germany; 1:100) overnight at 4°C followed by Alexa Fluor® 594-conjugated goat anti-mouse IgG secondary antibody (Molecular Probes, Eugene, OR, USA; 1:1000) for 1 hour at room temperature. Each antibody staining solution was 1x PBS containing 1% BSA. To visualize nuclei, cells were stained with 10 µg/ml Hoechst (Thermo Scientific Pierce) at room temperature for 5 minutes. Images were captured by Nikon Eclipse TE2000-E fluorescent microscopy equipped with MetaVue software (Molecular Devices, San Jose, CA, USA).

**Transwell migration assay**

Approximately $1\times10^5$ THP-1 cells were placed in the upper chamber of a 24-transwell plate (Corning Inc., Corning, NY, USA). Conditioned medium was prepared by diluting culture supernatants with an equal amount of fresh media plus 5% FCS. After 16h incubation, media was collected and cells counted by cellometer.

**Table S1: Genetically-engineered murine models of prostate cancer used in these studies**

| Mouse genotype (common name) | PB-Cre4 x PTEN$^{f/f}$ (Pten-deficient) | $ARR_2$PB-*MYC* (Hi-MYC) |
|---|---|---|
| Mouse background | C57Bl/6:129SvJ | FVB |
| Primary murine lobe | Anterior prostate | Ventral prostate |
| H&E histology | 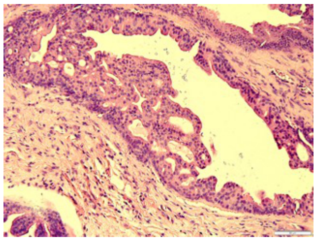 Intraductal histology infrequent in human PrCa | 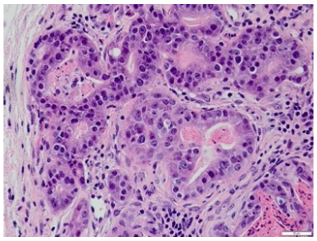 acinar histology most common human PrCa |
| Mouse age when tumor volume is ~300-500 mm$^3$ | ~ 6 months | ~ 10 months |

### Table S2: Rates of regression and recurrence for PTEN-deficient prostate cancers

#### Table S2a: Regression of PTEN-deficient prostate cancers post-castration

| Period monitored (post-CX) | Tumors imaged post-CX | Tumors that met regression criteria | Regression Rate |
|---|---|---|---|
| 6 wks | 18 | 14 | 78% |
| 10 wks | 41 | 36 | 88% |
| 16 wks | 53 | 46 | 87% |

PTEN-deficient prostate tumors were serially imaged by HFUS, and tumor volume was determined as per the Methods in the main text, for the indicated period of time (weeks) following castration

#### Table S2b: Recurrence of PTEN-deficient prostate cancers post-regression

| Period monitored (post-CX) | Tumors imaged after volume nadir | Tumors that met recurrence criteria | Recurrence Rate |
|---|---|---|---|
| 6 wks | 14 | 5 | 36% |
| 10 wks | 22 | 13 | 59% |
| 16 wks | 10 | 8 | 80% |

PTEN-deficient prostate tumors which met the RECIST-like criteria for regression were further serially imaged following the regression nadir by HFUS, and tumor volume was determined as per the Methods in the main text, for the indicated period of time (weeks) following castration

78% of prostate tumors in Pb-Cre4; $Pten^{fl/fl}$ mice showed reduced volume in response to castration by mouse-equivalent RECIST criteria within six weeks, when monitored using high-resolution, high-frequency ultrasound imaging (HFUS). This increased to 88% if HFUS monitoring for response was extended up to 10 weeks, but did not further increase (87% responsive) if monitored to 16 weeks after castration (Table S1a). After tumor volume nadir in response to castration, 36% (12/25) of tumors recurred by mouse-equivalent RECIST criteria within 6 weeks following castration as measured using HFUS imaging. Recurrence increased to nearly 60% by 10 weeks, and 80% (8/10) by 16 weeks after castration (Table S1b).

Mouse tumor volume equivalent of RECIST 1.1 therapy response is a single decline in overall volume of greater than 15.7% (equivalent to reduction of 30% in one axis on 2D imaging, the criterion for clinical partial response). Similarly, recurrence was determined by a sustained (two or more) increase in tumor volume of 10.5% from the nadir (equivalent to an increase of 20% in one axis, the criterion for progressive disease clinically). Reference: Morgan & Camidge (2108) *Reviewing RECIST in the era of prolonged and targeted therapy*. J. Thoracic Oncol. 13:154-164.

### Table S3: Statistical analysis of GEMM tumor regression and recurrence studies

Table S3A: Statistical analysis for Figure 4A

| | Regression | N | Prop. (95% CI) | Fisher's Test | Recurrence | N | Prop. (95% CI) | Fisher's Test |
|---|---|---|---|---|---|---|---|---|
| None | 4 | 4 | 1.00 (0.40-1.00) | Reference | 3 | 4 | 0.75 (0.19-0.99) | Reference |
| -3d | 0 | 5 | 0.00 (0.00-0.52) | p=0.0079 | NA | NA | NA | NA |
| -1d | 4 | 5 | 0.80 (0.28-0.99) | p=1.0000 | 0 | 4 | 0.00 (0.00-0.60) | p=0.1429 |
| +3d | 3 | 3 | 1.00 (0.29-1.00) | p=1.0000 | 2 | 3 | 0.67 (0.09-0.99) | p=1.0000 |

Table S3B: Statistical analysis for Figure 5C and 5D

| | PTEN-loss GEMM (Fig. 5C) | | | | Hi-MYC GEMM (Fig. 5D) | | | |
|---|---|---|---|---|---|---|---|---|
| | **Recurrence** | **N** | **Prop. (95% CI)** | **Fisher's Test** | **Recurrence** | **N** | **Prop. (95% CI)** | **Fisher's Test** |
| None/Vehicle | 3 | 4 | 0.75 (0.19-0.99) | Reference | 4 | 5 | 0.80 (0.28-0.99) | Reference |
| -1d | 0 | 6 | 0.00 (0.00-0.46) | p=0.0333 | 1 | 5 | 0.20 (0.01-0.72) | p=0.2063 |

Details: As per the Materials and Methods: for the small *n*, categorical data sets related to the studies using GEM prostate cancer models shown in Figures 4A, 5C and 5D, we employed the exact binomial test to determine the Clopper-Pearson confidence interval for the proportion and Fisher's exact test to determine the *p*-values. Note that the *p*-values reported from Fisher's exact test represent exact probabilities rather than asymptotic approximations. In data sets with small samples (such as treatments of the genetically-engineered mouse models of prostate cancer), these probabilities are discrete and should be interpreted cautiously, recognizing that conventional thresholds (e.g. $p$<0.05) may not directly apply.

### FIGURE S1

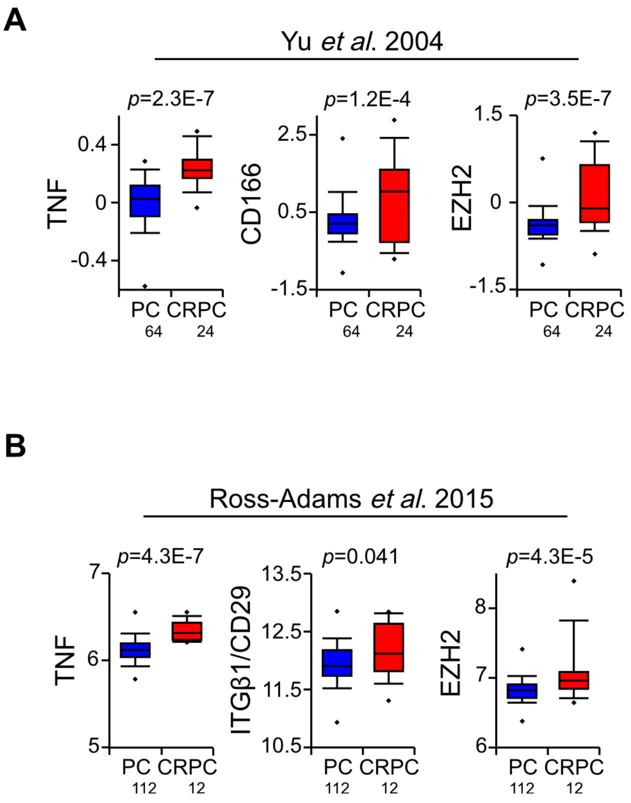


**Supplementary Figure S1. Expression of basal/stem cell related genes and TNF in human prostate cancer samples.** **A.** TNF and stem cell markers mRNA expression in human primary PC (blue) and CRPC (red) from the indicated publications, extracted and analyzed at the Oncomine.org (**A**) or GEO (**B**) web browsers, and plotted as box and whisker plots. Sample size (bottom) and p-value (top) are indicated. PC and CRPC sample sets were compared by Student's unpaired *t*-test. We previously reported the TNF expression analysis (in the left-most part of panel A) of the Yu et al. 2004 dataset in a separate set of correlative gene expression analyses (10).

### FIGURE S2 - Extended data for Figure 2 (adds LNCaP cell line)

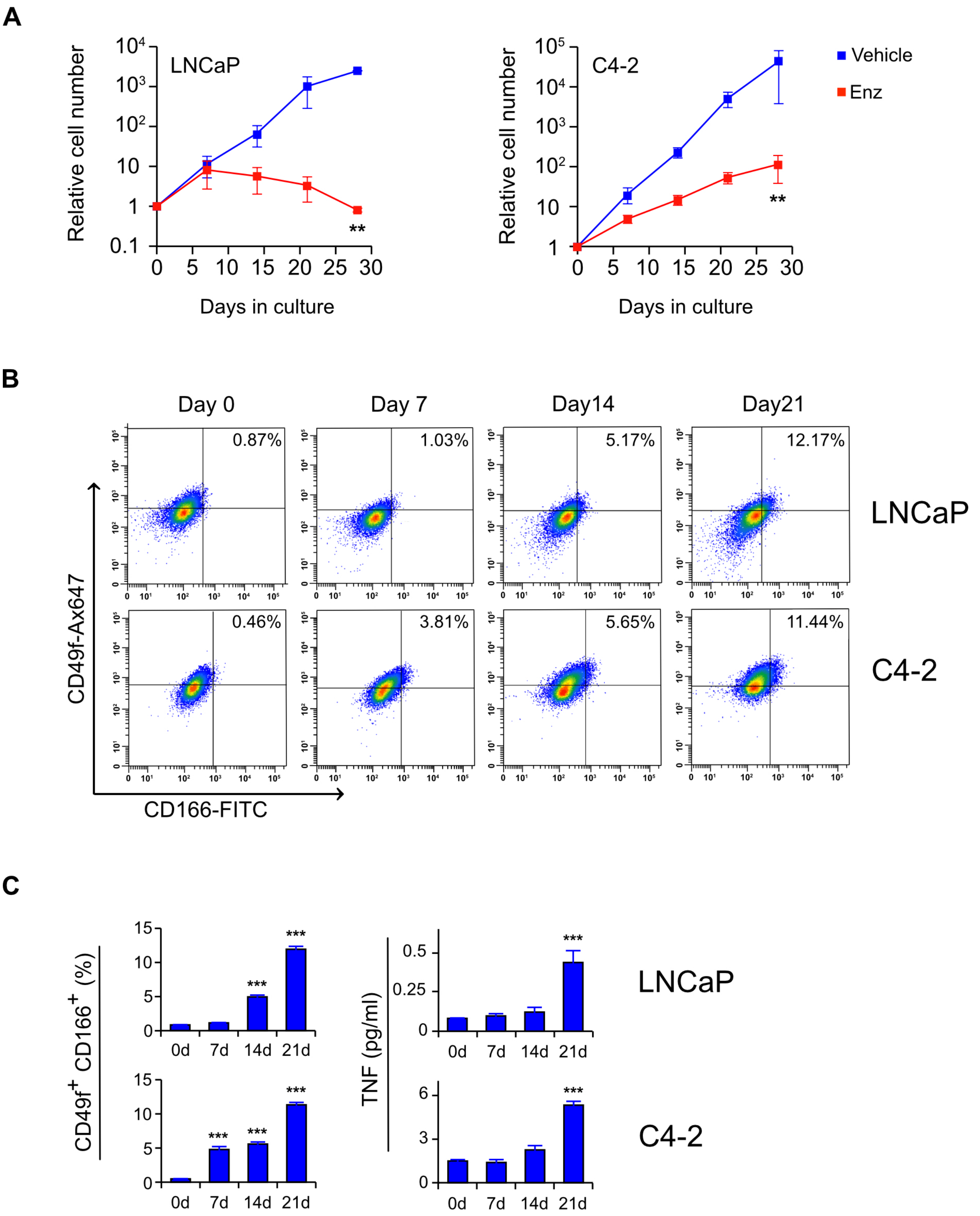

### LEGEND CORRESPONDING TO THE FIGURE ON THE PREVIOUS PAGE:

**Supplementary Figure S2. Extended enzalutamide treatment selects for basal stemness and TNF secretion: extended version of Figure 2.** This figure is an expanded version of Figure 2, with reproduction of the C4-2 data from Figure 2, plus additional results from similar experiments using LNCaP, a more androgen-dependent cell line that is the parental cell line for C4-2. C4-2 or LNCaP cells were grown in media plus 10% serum and treated with vehicle (Vehicle; blue) or 10 µM enzalutamide (Enz; red) for the indicated time. **A.** LNCaP (left) and C4-2 (right) cell growth curve in the presence of enzalutamide. Cells were counted microscopically at the indicated times. Data are shown as mean ± s.e.m. (n=3). **B.** FACS analysis for basal cell stemness markers in enzalutamide treated LNCaP (upper row) and C4-2 (lower row) cells. Cells grown in enzalutamide for the indicated time were incubated with the indicated fluor-labeled antibodies and analyzed by FACS. The relative fluorescence intensity is represented as dot plots. The fraction of the cells that correspond to the $CD49f^{hi}/CD166^{hi}$ population (upper, right quadrant) is indicated. **C.** TNF secretion and corresponding Fraction of cells scoring $CD49f^{hi}/CD166^{hi}$ and corresponding TNF secretion. Data from FACS analyses (panel **B**) is plotted on the left and TNF determined by ELISA is plotted on the right. Values are given as mean ± s.e.m. ($n$ = 2). **, $p$ < 0.01 ***$p$ < 0.001 (two-way ANOVA followed by Tukey-Kramer HSD test).

### LEGEND CORRESPONDING TO FIGURE ON THE FOLLOWING PAGE:

**Supplementary Figure S3. Extended enzalutamide treatment selects for basal stemness and TNF secretion: immunofluorescence staining of basal markers CD49f and CD166.** LNCaP and C4-2 cells were treated with vehicle (–, blue) or 10 µM enzalutamide (Enz; +, red) for the indicated time. Aliquots of the corresponding cultures were incubated with either anti-CD49f or anti-CD166 antibodies, followed by appropriate fluorescent-tagged secondary antibodies. Hoechst stain was used to identify nuclei. Representative photomicrographs were captured under appropriate fluorescence and fluorescent cells counted. **Panels A-D.** LNCaP cell line. Fluorescent microscopic images corresponding to CD49f (**A**) and CD166 (**D**, left side) following the indicated treatments. The corresponding quantitation is plotted for CD49f (**B**) and CD166 (**D**, right side). The TNF levels, determined by ELISA of the media for the indicated cultures (**C**). Plotted values are given as mean ± s.e.m. of percentage of positively stained cells ($n$ = 3). ***$p$<0.001(Two-way ANOVA followed by Tukey-Kramer HSD test). **Panels E-H.** C4-2 cell line. Analogous to those shown for LNCaP in panels **A-D**.

**FIGURE S3**

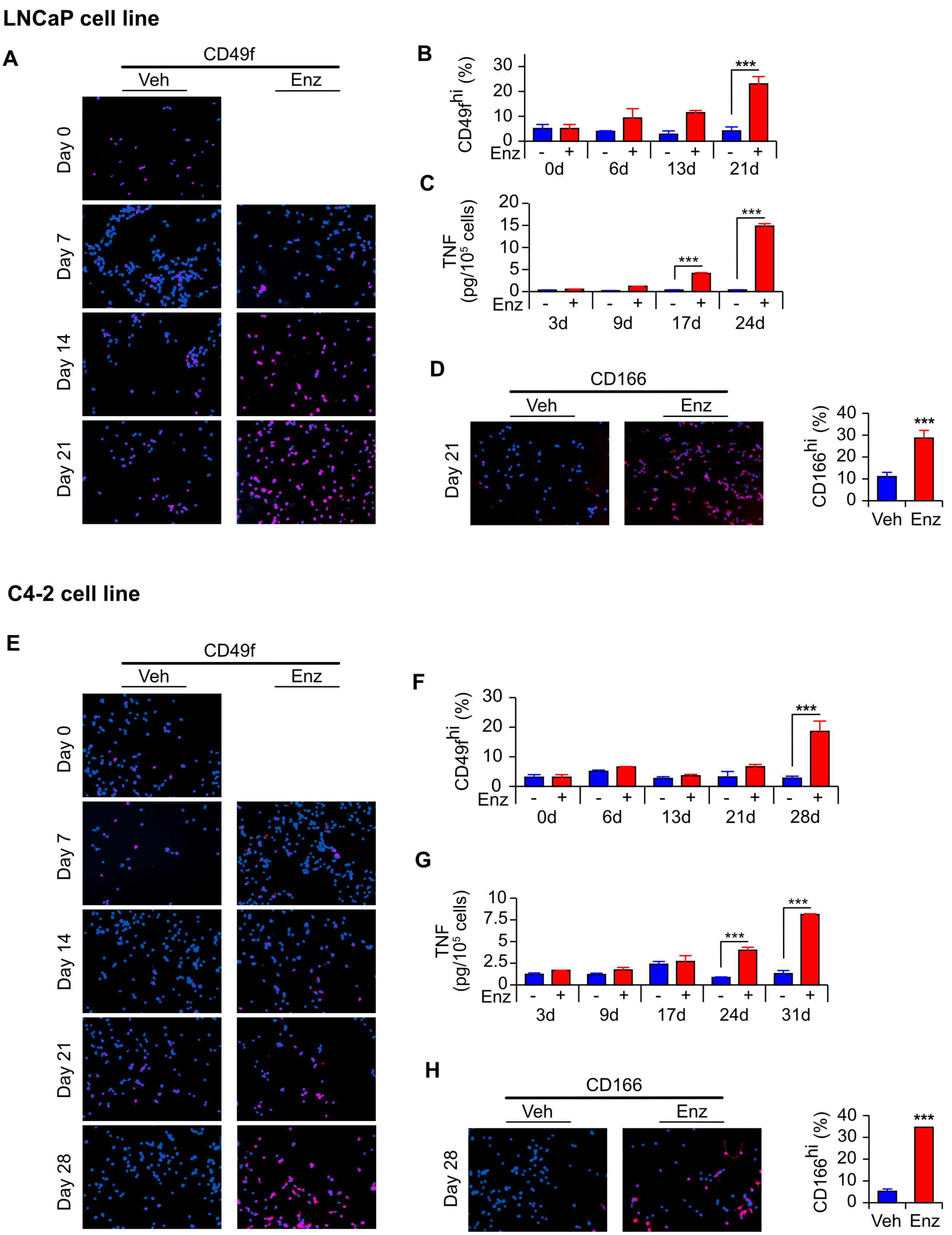
LNCaP cell line
A
CD49f
Veh
Enz
Day 0
Day 7
Day 14
Day 21
B
CD49f^hi (%)
30
20
10
0
Enz
- + - + - + - +
0d 6d 13d 21d
***
C
TNF (pg/10^5 cells)
20
15
10
5
0
Enz
3d 9d 17d 24d
***
***
D
CD166
Veh
Enz
Day 21
CD166^hi (%)
40
30
20
10
0
Veh Enz
***
C4-2 cell line
E
CD49f
Veh
Enz
Day 0
Day 7
Day 14
Day 21
Day 28
F
CD49f^hi (%)
30
20
10
0
Enz
0d 6d 13d 21d 28d
***
G
TNF (pg/10^5 cells)
10
7.5
5
2.5
0
Enz
3d 9d 17d 24d 31d
***
***
H
CD166
Veh
Enz
Day 28
CD166^hi (%)
40
30
20
10
0
Veh Enz
***

### FIGURE S4

#### A: Autocrine CCL2: Pr ca cell line mono-cultures

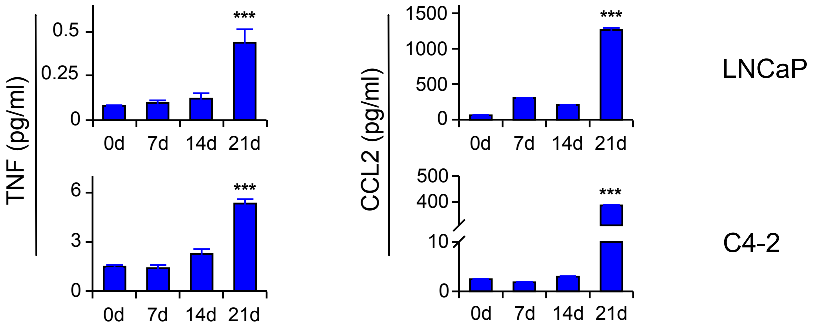


#### B: Autocrine CCL2: ENZ-induced CCL2 production is inhibited by anti-TNF

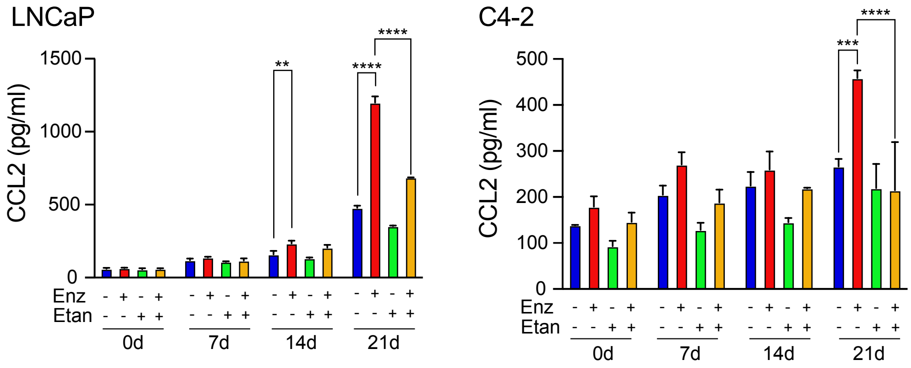


#### C: Paracrine CCL2: Pr ca cell line + WPMY-1 myofibroblast-like cell line co-cultures

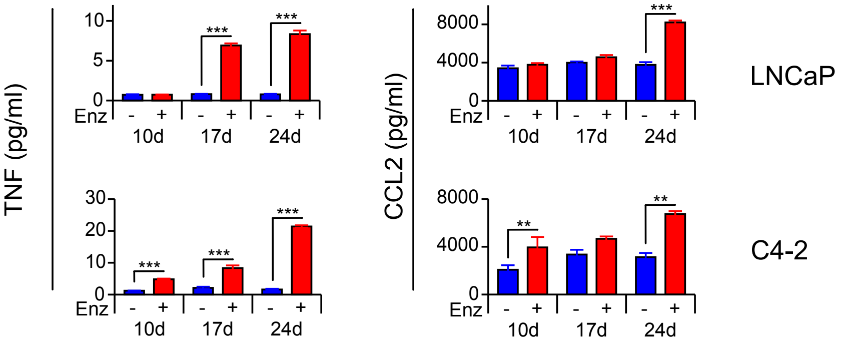


#### D: CCR2$^+$ THP-1 (macrophage-like cells) transwell migration

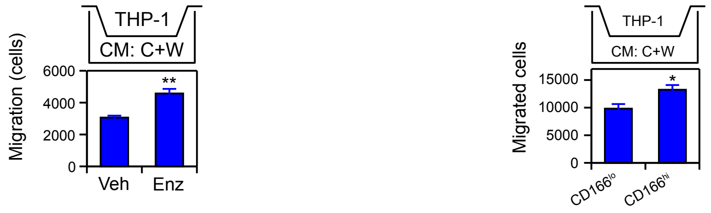


Conditioned media from a mono-culture of:
C4-2 cells treated with vehicle or enz

Conditioned media from a co-culture of:
CD166 sorted C4-2 cells & WPMY-1 cells

**Supplementary Figure S4. Enzalutamide induces CCL2 secretion under autocrine and paracrine conditions. A.** Autocrine CCL2 production by LNCaP and C4-2 mono-cultures treated with enzalutamide for the indicated times. The TNF ELISA data is reproduced from Figure 2C. The CCL2 ELISA data is from the same cultures**. B.** Similar to A, except that some cultures were treated with vehicle or etanercept, as indicated. **C.** Paracrine CCL2 production by prostate cancer cell lines (LNCaP and C4-2) co-cultured with WPMY-1 myofibroblasts. TNF and CCL2 were assayed by ELISA using the media from cultures treated as indicated. For panels **A-C**, values are given as mean ± s.e.m. ($n$ = 2). **, $p$ < 0.01 ***$p$ < 0.001 (Two-way ANOVA followed by Tukey-Kramer HSD test). **D.** Migration of THP-1 macrophage-like cells towards conditioned media from enzalutamide treated co-cultures. As diagrammed, transwell analysis of THP-1 migration, toward conditioned media from C4-2/WPMY-1 co-cultures (CM: C+W). Left: co-cultures were treated as indicated. Right: Live C4-2 cells sorted into CD166$^{lo}$ and CD166$^{hi}$ fractions as described in Figure 3A, then separately co-cultured with WPMY-1 cells. Data are given as mean ± s.e.m. ($n$ = 3), *p<0.05, **$p$<0.01, ***$p$<0.001 (Student's unpaired $t$-test).

### FIGURE S5

**A**

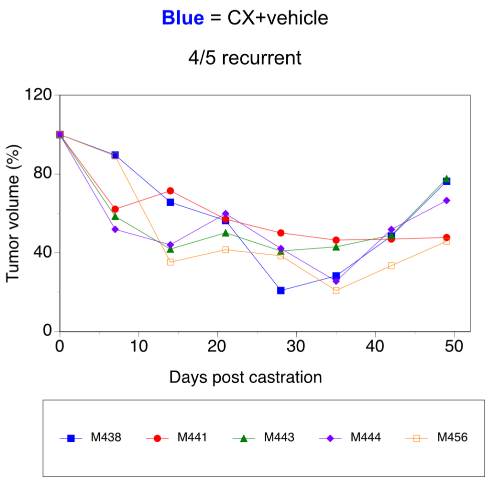


**B**

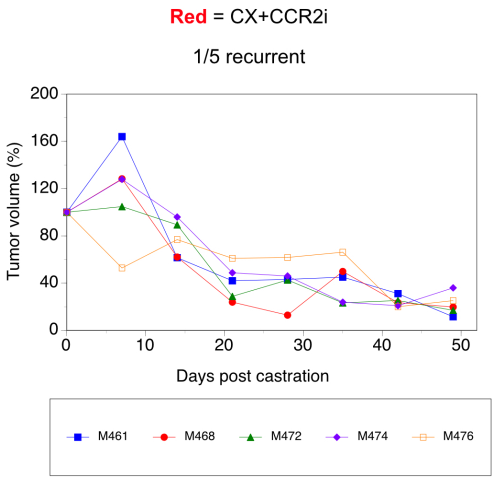


**Supplementary Figure S5. CCR2 inhibitor suppresses castration-induced regression in Hi-MYC prostate cancer-bearing mice: related to Figure 5D.** Tumor volume kinetics for individual mice that are summarized/averaged in Figure 5D.

### FIGURE S6

**A**

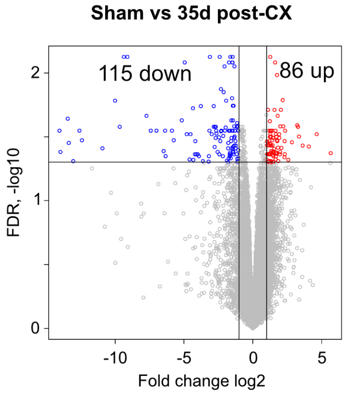


**B**

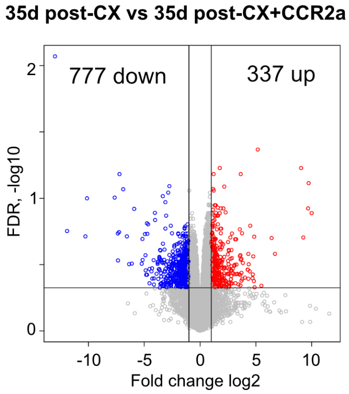


**Supplementary Figure S6. Volcano plots of differentially-expressed gene (DEG) analysis: related to Figure 6.** RNA was extracted from PTEN-deficient tumors from the following mice: sham-operated (Sham), 35d post-castration (35d post-CX) and 35d post-castration after receiving CCR2a (35dp-CX+CCR2a). DEG was performed on bulk RNAseq data using RNAseq transcript sets derived from tumors in mice that were sham-operated vs mice 35d post-castration (**A**) or from tumors in mice 35d post-castration vs mice treated with CCR2a 35d post-castration (**B**). Select genes with FDR<0.05 and |log2FC| up-regulated in the bulk tumor mRNA from mice 35 days post-castration versus sham-operated mice (**A**, upper right quadrant) or down-regulated in the bulk tumor mRNA from mice 35 days post-castration versus those treated with CCR2a (**B**, upper left quadrant) were subjected to GO analysis (Fig. 6).

### FIGURE S7

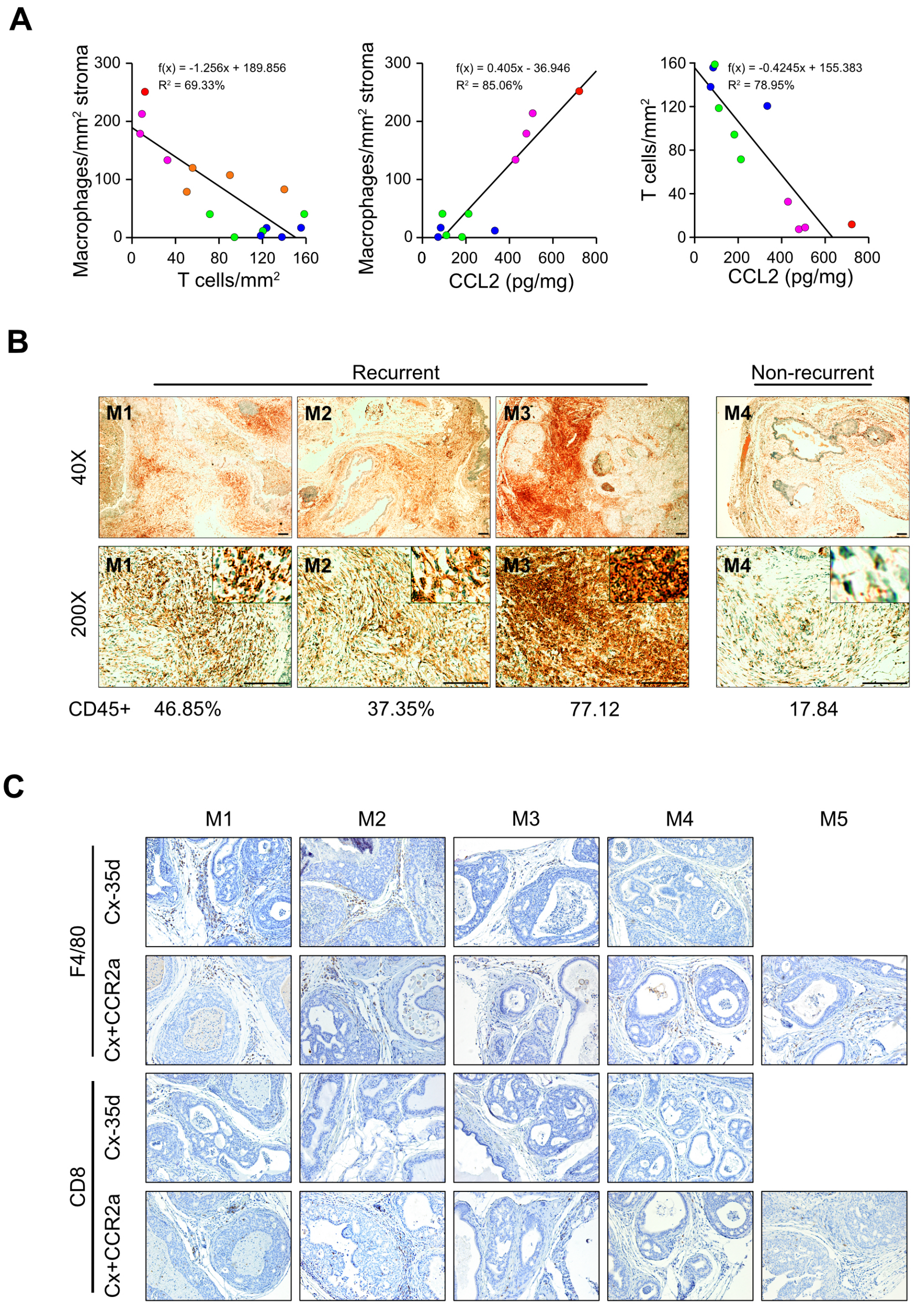


**Supplementary Figure S7. Supplementary data in support of castration-induced immunosuppression**. **A.** TAM density, CD8 T cell density and CCL2 are tightly correlated. There is an inverse correlation between TAMs and T cells, a positive correlation between TAMs and CCL2 levels and an inverse correlation between T cells and CCL2. Blue=sham, green=14d post-castration, magenta=35d post-castration from recurrent tumors, red=35d post-castration from non-recurrent tumors. **B.** TAMs in recurrent and non-recurrent tumors, harvested 35d post-castration. IHC-staining (red) for F4/80$^+$ macrophages in recurrent and non-recurrent tumors, as indicated. Percentage of CD45+ immune cells in the tumor section are indicated at bottom, for each mouse. Green counter-staining was employed. Scale bar = 500 μm (40X), 100 μm (200X). **C.** Additional data for Figure 7B. IHC-stained sections for all mouse tumors (only tumor M1 is shown in Fig. 7B), all photographed at 40X magnification.

### FIGURE S8

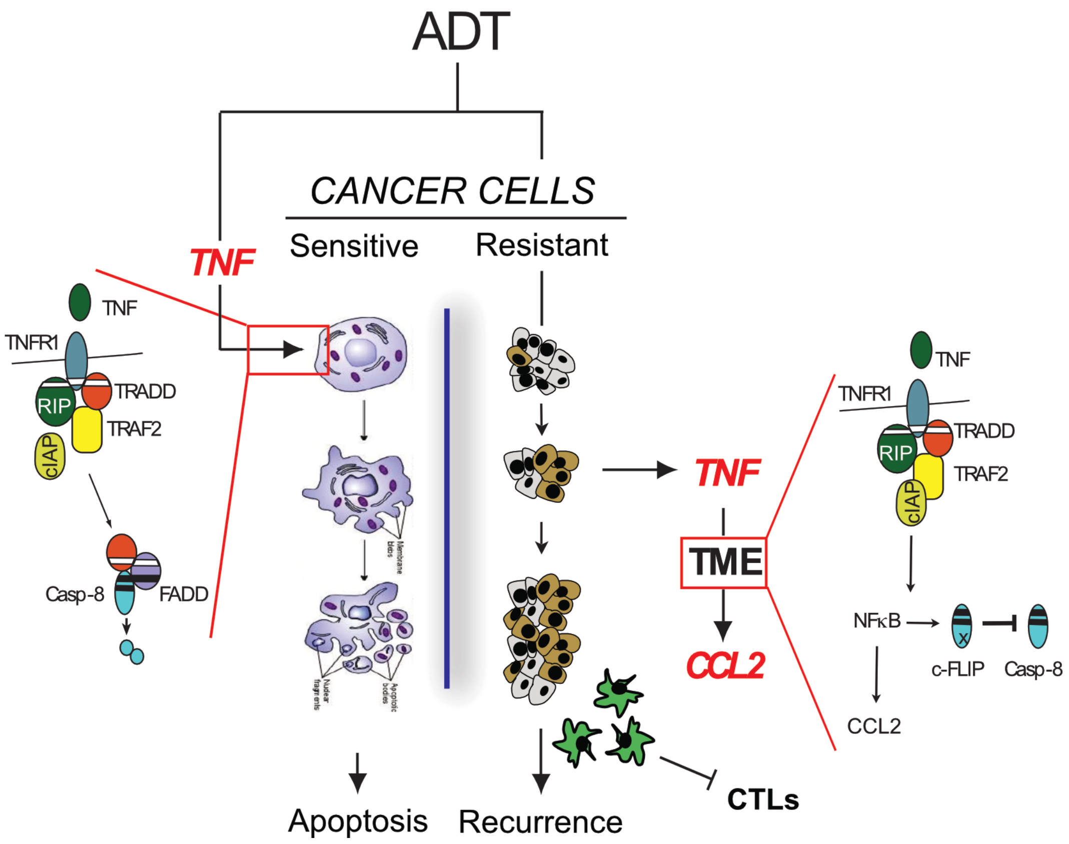


**Supplementary Figure S8. TNF signaling switch drives regression and recurrence of prostate cancers following androgen deprivation (castration).** The cartoon illustrates the two opposing TNF signaling responses (anti-tumor versus pro-tumor) of murine prostate cancers to castration/ androgen deprivation therapy. **LEFT:** About a week post-castration, pre-existing TNF in the TME binds TNFR1 on the prostate epithelial cell surface to trigger casapase-8 activation/ cleavage resulting in prostate epithelial cell apoptosis, as first described by Kerr *et al.* (53). This leads to tumor regression. Note that we also recently described the very early (within 1d of castration) onset of 'vascular regression' (8), which is also TNF-dependent. The likely mechanism of vascular regression is TNF-induced apoptosis of endothelial cells, inducing transient hypoxia and perhaps p53-mediated stress that could contribute to regression. **RIGHT:** Following regression due to apoptosis of tumor epithelium (light grey cytoplasm) there is a relative increase in cells with a basal stem-cell phenotype (dark brown cytoplasm) which are the source of a late surge in TNF secretion into the TME. We propose that the relatively high TNF levels induce the expression of NFκB in multiple tumor-associated cell populations. High NFκB in epithelial tumor cells blocks caspase-8 activation by inducing transcription of the gene encoding c-FLIP, an inactive homologue of caspase-8 that acts as a dominant-negative inhibitor. More importantly, NFκB also induces CCL2 gene transcription in macrophages and perhaps other TME cells, acts to recruit TAMs (green cells) to the TME and promote immunosuppression, and reduces the number or anti-tumor effectiveness of cytotoxic tumor lymphocytes (CTLs).
